# *In Vivo* Neuroregeneration to Treat Ischemic Stroke in Adult Non-Human Primate Brains through NeuroD1 AAV-based Gene Therapy

**DOI:** 10.1101/816066

**Authors:** Long-Jiao Ge, Fu-Han Yang, Jie Feng, Nan-Hui Chen, Min Jiang, Jian-Hong Wang, Xin-Tian Hu, Gong Chen

**Affiliations:** Key Laboratory of Animal Models and Human Disease Mechanisms of Chinese Academy of Sciences & Yunnan Province, Kunming Institute of Zoology, Chinese Academy of Sciences, Kunming, Yunnan 650223, China; Guangdong-Hongkong-Macau Institute of CNS Regeneration, Jinan University, Guangzhou 510632, China; Department of Biology, Huck Institutes of Life Sciences, Pennsylvania State University, University Park, PA 16802, USA; State Key Laboratory of Medical Neurobiology and MOE Frontier Center for Brain Science, Institutes of Brain Science, Fudan University, Shanghai 200032, China; CAS Center for Excellence in Brain Science, Chinese Academy of Sciences, Shanghai, 200031, China; Kunming Primates Research Center, Kunming Institute of Zoology, Chinese Academy of Sciences, Kunming, Yunnan650223, China; Kunming College of Life Science, University of the Chinese Academy of Sciences, Kunming, Yunnan 650204, China

**Keywords:** Reprogramming, in vivo, cell conversion, astrocyte, neuron, non-human primate, stroke, ischemic injury, brain repair, neuroregeneration

## Abstract

Stroke is a leading cause of death and disability but most of the clinical trials have failed in the past, despite our increasing understanding of the molecular and pathological mechanisms underlying stroke. While many signaling pathways have been identified in the aftermath of stroke, the majority of current approaches are focusing on neural protection rather than neuroregeneration. In this study, we report an *in vivo* neural regeneration approach to convert brain internal reactive astrocytes into neurons through ectopic expression of a neural transcription factor NeuroD1 in adult non-human primate (NHP) brains following ischemic stroke. We demonstrate that NeuroD1 AAV-based gene therapy can convert reactive astrocytes into neurons with high efficiency (90%), but astrocytes are never depleted in the NeuroD1-expressed areas, consistent with the proliferative capability of astrocytes. The NeuroD1-mediated *in vivo* astrocyte-to-neuron (AtN) conversion in monkey cortex following ischemic stroke increased local neuronal density, reduced reactive microglia, and surprisingly protected parvalbumin interneurons in the converted areas. The NeuroD1 gene therapy showed a broad time window, from 10 days to 30 days following ischemic stroke, in terms of exerting its neuroregenerative and neuroprotective effects. The cortical astrocyte-converted neurons also showed Tbr1+ cortical neuron identity, similar to our earlier findings in rodent animal models. Unexpectedly, NeuroD1 expression in converted neurons showed a significant decrease after 6 months of viral infection, suggesting a potential self-regulatory mechanism of NeuroD1 in adult mature neurons of NHPs. These results suggest that *in vivo* cell conversion through NeuroD1-based gene therapy may be an effective approach to regenerate new neurons in adult primate brains for tissue repair.

## INTRODUCTION

Stroke is a leading cause of serious long-term disability and mortality worldwide, accounting for 11.8% of total death (Feigin et al., 2014). Stroke prevalence in adults is 2.7% in the United States, with ∼800,000 people suffering from a stroke every year (Benjamin et al., 2018). Given the rapidly aging population and the risk of stroke increasing significantly in senior people more than 60 years old, the number of people with stroke will continuously increase (Ovbiagele et al., 2013). Ischemic stroke is accounting for ∼87% of all strokes, and current treatments mainly target on the re-establishment of blood flow and neuroprotection (Avasarala 2015). Stroke survivors often experience long-term disabilities due to a substantial loss of neuronal cells caused by ischemic injury. Therefore, in order to achieve better functional recovery after stroke, it is pivotal to regenerate new neurons after stroke to replenish the lost neurons and restore the lost brain functions.

Our group previously reported that *in vivo* overexpression of a single neural transcription factor NeuroD1 can convert endogenous glial cells directly into functional neurons (Guo et al., 2014), providing a new approach for neural repair in the brain and spinal cord. Our more recent studies also demonstrated that NeuroD1 AAV-based gene therapy has great potential in repairing the ischemic injured or stab lesioned mouse cortex through direct *in vivo* astrocyte-to-neuron conversion (Chen et al., 2019; Zhang et al., 2018). Besides NeuroD1, other groups have reported that expression of transcription factors Ngn2, Ascl1, Sox2, or combinations of factors can also convert internal glial cells into neurons in the brain or spinal cord (Gascon et al., 2016; Grande et al., 2013; Jorstad et al., 2017; Liu et al., 2015; Niu et al., 2015; Niu et al., 2013; Pereira et al., 2017; Rivetti di Val Cervo et al., 2017; Su et al., 2014). In addition to the overexpression of transcription factors, we have also demonstrated that small molecules can directly convert human glial cells into functional neurons (Yin et al., 2019; Zhang et al., 2015) through transcriptome-wide activation of neuronal genes (Ma et al., 2019). Other groups also reported direct chemical reprogramming of glial cells (Gao et al., 2017) or fibroblast cells (Ambasudhan et al., 2011; Hu et al., 2015; Li et al., 2015) into neurons. These studies point to the possibility of using brain internal glial cells to directly generate new neurons for brain repair. However, in order to test the therapeutic potential of *in vivo* cell conversion technology in human clinical trials, it is important to first understand in non-human primate (NHP) models besides rodent studies, because mouse brain is very different from human brain in terms of volume, structure, and genetic compositions. Many experimental parameters obtained from rodent animal models cannot be extrapolated to human patients, evidenced by many failed stroke clinical trials over the past decades (Diener et al., 2008; O’Collins et al., 2006; Percie du Sert et al., 2017; Turner et al., 2013).

Here, we provide direct evidence in Rhesus Macaque monkeys that expressing a single neural transcription factor NeuroD1 in reactive astrocytes caused by ischemic injury can convert them into neurons at the injury site. Following the *in situ* astrocyte-to-neuron (AtN) conversion, the neuronal density in the NeuroD1-treated injury areas were significantly increased, accompanied by an increase of dendritic marker and synaptic marker. Unexpectedly, parvalbumin interneurons were significantly protected from ischemic injury in NeuroD1-infected areas, while microglia was significantly reduced after NeuroD1-treatment. Our findings suggest that *in vivo* cell conversion technology not only regenerate new neurons in the injury areas, but also ameliorate the microenvironment to be more neuro-protective and neuro-permissive. These studies in NHPs make one step closer toward future therapeutic treatment for ischemic stroke and possibly for other neurological disorders as well.

## RESULTS

### NeuroD1-mediated astrocyte-to-neuron conversion in the monkey cortex

We have previously demonstrated that reactive astrocytes inside mouse brains can be directly converted into functional neurons through expressing a single neural transcription factor NeuroD1 (Chen et al., 2019; Guo et al., 2014; Zhang et al., 2018). Because mouse brains are far different from human brains, it is uncertain whether such *in vivo* cell conversion technology would be applicable for future human clinical trials. To overcome such uncertainty, we decided to further investigate whether such *in vivo* neuroregeneration approach can be reproduced successfully in adult NHP brains in order to pave the way towards future clinical therapies.

Adult Rhesus Macaque monkeys (male) aged from 9 to 21 years old were used in this study. AAV-based gene therapy was employed to test the efficacy of NeuroD1 (Chen et al., 2019) in converting the reactive astrocytes following ischemic cortical stroke into neurons in the monkey cortex. To infect the reactive astrocytes, we used astrocytic promoter GFAP to drive the expression of NeuroD1 or control reporter GFP. We first injected the control AAV9 GFAP::GFP viruses into the cortex of Rhesus Macaque monkey (monkey ID #04339) and found that the majority of GFP-infected cells were GFAP+ astrocytes at 28 days post viral infection (dpi), as expected (Fig. 1A). In contrast, when injecting AAV9 GFAP::NeuroD1-GFP into the monkey cortex, we found that many NeuroD1-GFP-infected cells (both GFP+ and NeuroD1+) became NeuN+ neurons at 28 dpi (Fig. 1B). Some NeuroD1-converted neurons also showed neuronal dendritic marker MAP2 (Fig. 1C). Interestingly, like that reported in rodent models (Chen et al., 2019; Zhang et al., 2018), we found that some NeuroD1-GFP infected cells were immunopositive for both GFAP (red) and NeuN (blue) (Fig. 1D), suggesting a transitional stage in-between astrocytes and neurons during the conversion process. Quantified data found that 99% of GFAP::GFP infected cells were GFAP+ astrocytes, but 46% of NeuroD1-GFP infected cells already became NeuN+ neurons within one month of infection (Fig. 1E-F). Among all the NeuroD1-GFP infected cells at 28 dpi, besides 46% of NeuN+ neurons, there were also 35% GFAP+ astrocytes and 7% of NeuN+/GFAP+ cells together with 22% NeuN-/GFAP-cells (Fig. 1G). It appears that both NeuN+/GFAP+ double positive cells and NeuN-/GFAP-double negative cells were transitional cells during AtN conversion process. These results suggest that overexpression of NeuroD1 in the astrocytes of monkey brains can convert them into neurons, similar to that found in the mouse brains (Chen et al., 2019; Guo et al., 2014; Zhang et al., 2018).

**Figure 1.**
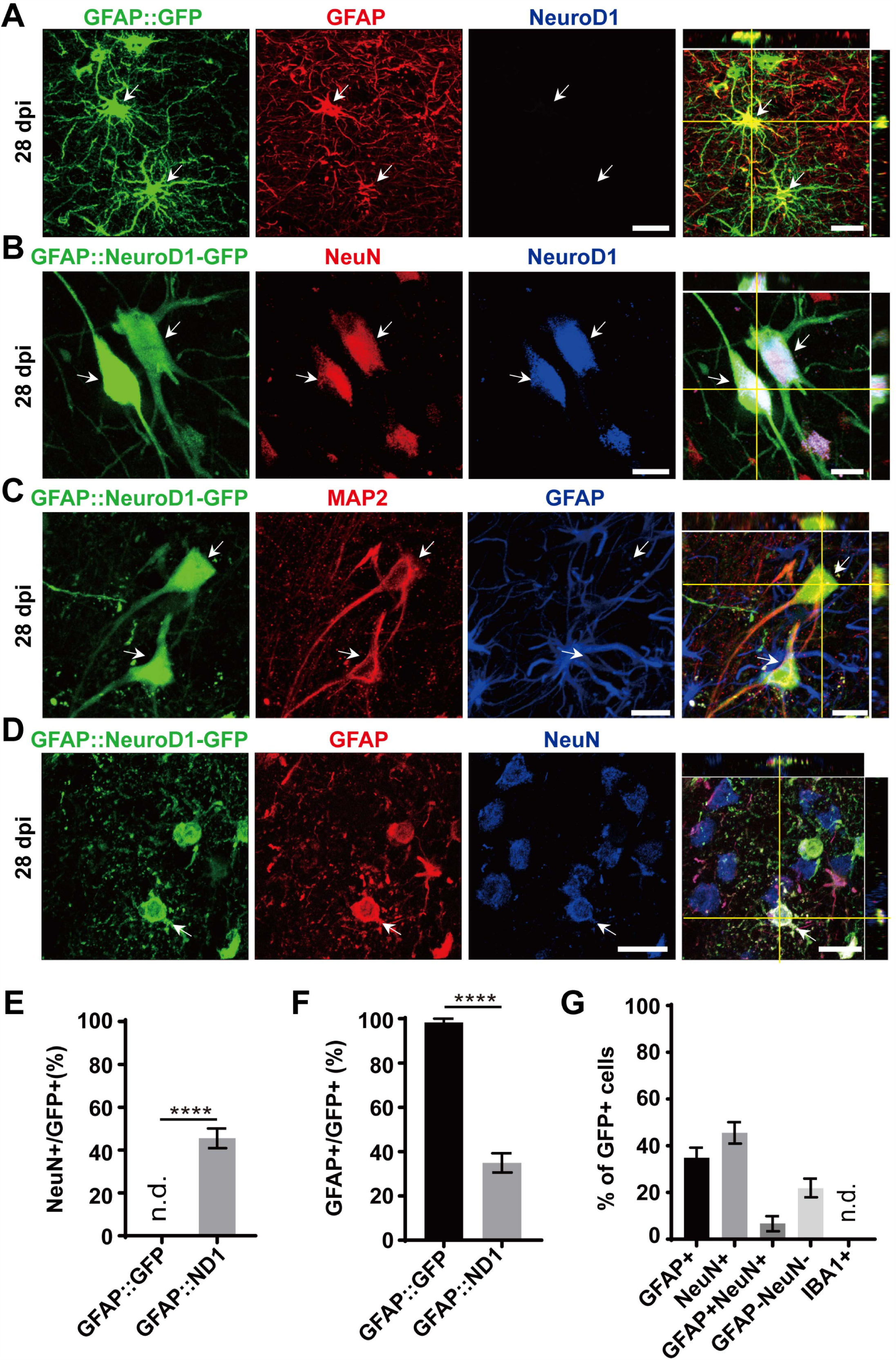
NeuroD1-mediated astrocytes-to-neuron conversion in the monkey cortex. **(A)** Injection of AAV9 viruses expressing GFP alone under human GFAP promoter (hGFAP::GFP) into monkey cerebral cortex infected astrocytes only with colocalization of astroglial marker GFAP (red) at 28 days post viral injection (dpi). These cells were negative for NeuroD1 staining. Scale bar, 20 μm. **(B-C)** Injection of AAV9 viruses expressing NeuroD1-GFP under human GFAP promoter (hGFAP::NeuroD1-P2A-GFP) converted astrocytes into neurons with colocalization of neuronal marker NeuN (B) and dendritic marker MAP2 (C). The converted neurons (28 dpi) were confirmed to express NeuroD1 (B) but lost GFAP signal (C). Scale bars, 20 μm. **(D)** Some hGFAP::NeuroD1-P2A-GFP infected cells (green) were immunopositive for both NeuN and GFAP, suggesting a transitional stage in-between astrocytes and neurons during the conversion process. Scale bar, 20 μm. **(E-F)** Cell counting analysis of NeuN+ or GFAP+ cells among viral infected cells. Data are presented as mean ± SEM. N = 30 random fields from triplicate samples in the control group; N = 60 random fields from triplicate samples in the ND1 group. n.d., not detected. ****p < 0.0001, unpaired two-tailed Student’s *t*-test. **(G)** Bar graphs showing the different cell type compositions among all the NeuroD1-GFP infected cells. Data presented as mean ± SEM. N = 30 random fields from triplicate samples. n.d., not detected.

While GFAP promoter-driven NeuroD1-GFP expression converted astrocytes into neurons, some converted neurons might have lost GFP signal due to the reduced GFAP promoter activity during the neuronal conversion process, underestimating the actual conversion efficiency. To overcome this problem, we developed a Cre-FLEX system to separate GFAP promoter from NeuroD1 expression in two different AAV vectors (Chen et al., 2019). Specifically, Cre recombinase was controlled by GFAP promoter to target its expression in astrocytes (GFAP::Cre), while NeuroD1 expression was controlled by CAG promoter and flanked by LoxP sites in an inverted fashion (FLEX-CAG::NeuroD1-P2A-GFP or mCherry). To test the specificity of this Cre-FLEX system, we injected GFAP::GFP together with GFAP::Cre and FLEX-CAG::mCherry-P2A-mCherry or FLEX-CAG::NeuroD1-P2A-mCherry into the monkey cortex (Fig. 2A). As expected, GFAP::GFP-infected cells were all GFAP-positive astrocytes (Fig. 2B). Similarly, GFAP::Cre and FLEX-CAG::mCherry-P2A-mCherry also infected GFAP+ astrocytes, same as those infected by GFAP::GFP (Fig. 2B). In contrast, when GFAP::GFP were injected together with GFAP::Cre and FLEX-CAG::NeuroD1-P2A-mCherry, many of the GFP and mCherry co-infected cells became NeuN+ neurons, suggesting successful conversion of astrocytes into neurons after NeuroD1 expression (Fig. 2C, white arrowhead). Some NeuroD1-mCherry converted neurons showed much weaker GFP signal (Fig. 2C, dashed circle), suggesting that the GFAP promoter activity was downregulated in the newly converted neurons which led to reduced GFP expression for GFAP::GFP viral vector. Note that the cells infected only by GFAP::GFP (no NeuroD1-mCherry) were still astrocytes (Fig. 2D, green arrow), even though the neighboring astrocytes co-infected with NeuroD1-mCherry had become neurons (Fig. 2D, white arrowhead). Quantitation of the NeuN+ cells among the viral infected cells by both GFAP::GFP and GFAP::Cre plus Flex-NeuroD1-mCherry revealed high neuronal conversion efficiency (94.4 ± 5.5%) in monkey cortex. Together, these results demonstrate that AAV NeuroD1 Cre-FLEX system can successfully convert monkey cortical astrocytes into neurons and that the astrocyte-converted neurons can be labeled with reporters for long-term investigation.

**Figure 2.**
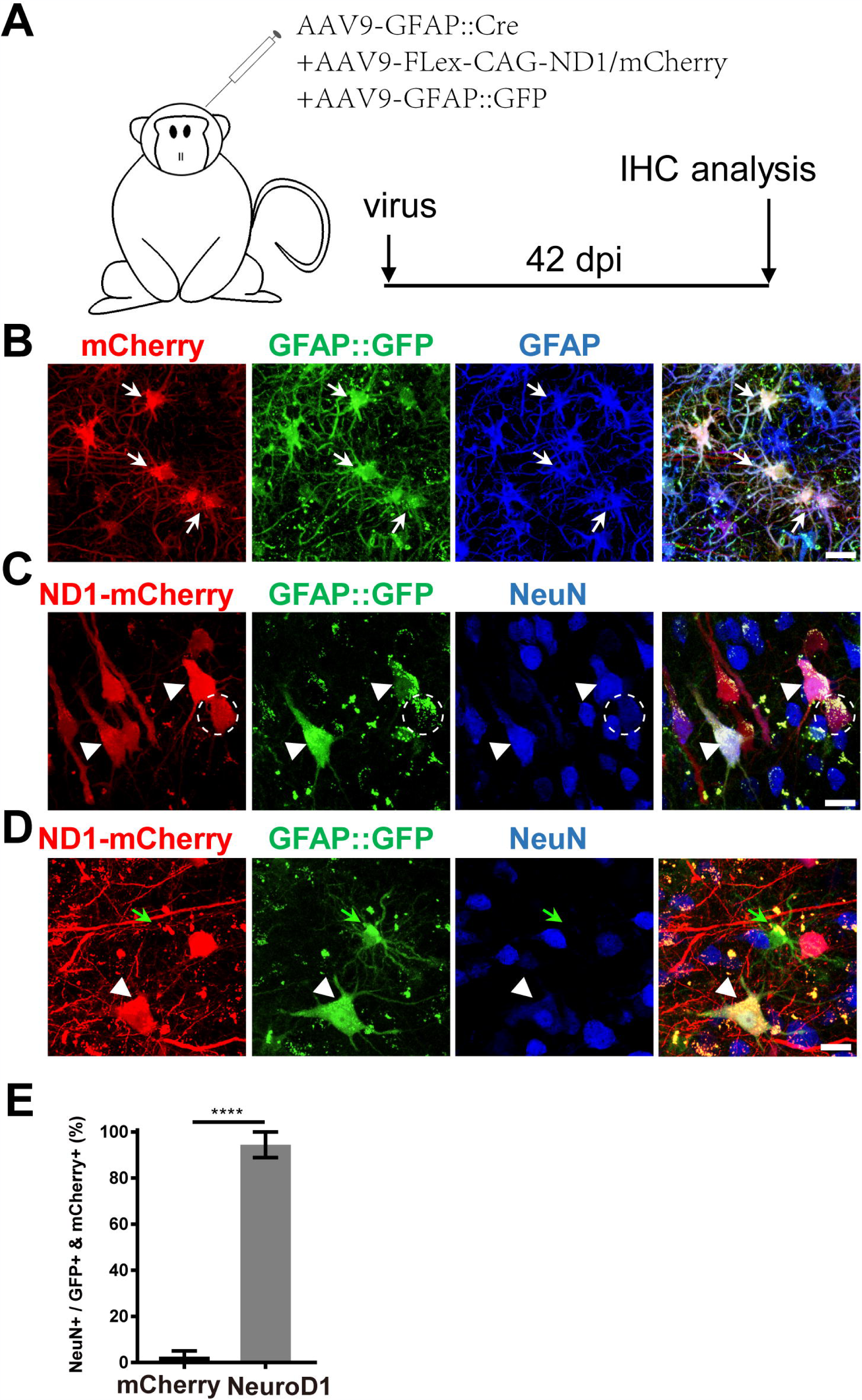
An engineered Cre-Flex system for high efficiency conversion in the monkey cortex. **(A)** Monkey cortex was injected with AAV9 hGFAP::GFP + GFAP::Cre together with either Flex-CAG::mCherry (control) or Flex-CAG::NeuroD1-mCherry and analyzed at 42 dpi. AAV9 GFAP::GFP was used to label local astrocytes in the cortex. **(B)** Representative images showing astrocytes (GFAP+) co-infected by the control viruses (AAV9 GFAP::GFP + GFAP::Cre + Flex-CAG::mCherry). Scar bars, 20 μm. **(C-D)** Representative images showing converted neurons (arrowhead, NeuN+) after infected by NeuroD1 viruses (AAV9 GFAP::GFP + GFAP::Cre + Flex-CAG::NeuroD1-mCherry) at 42 dpi. Note that GFAP::GFP-only infected cells were still astrocytes (D, green arrow). Scar bars, 20 μm. **(E)** Quantification of the neuronal conversion efficiency (NeuN+/GFP+mCherry+) among mCherry control (2.5 ± 2.5%) and NeuroD1-mCherry infected cells (94.4 ± 5.5%).

**Figure 3.**
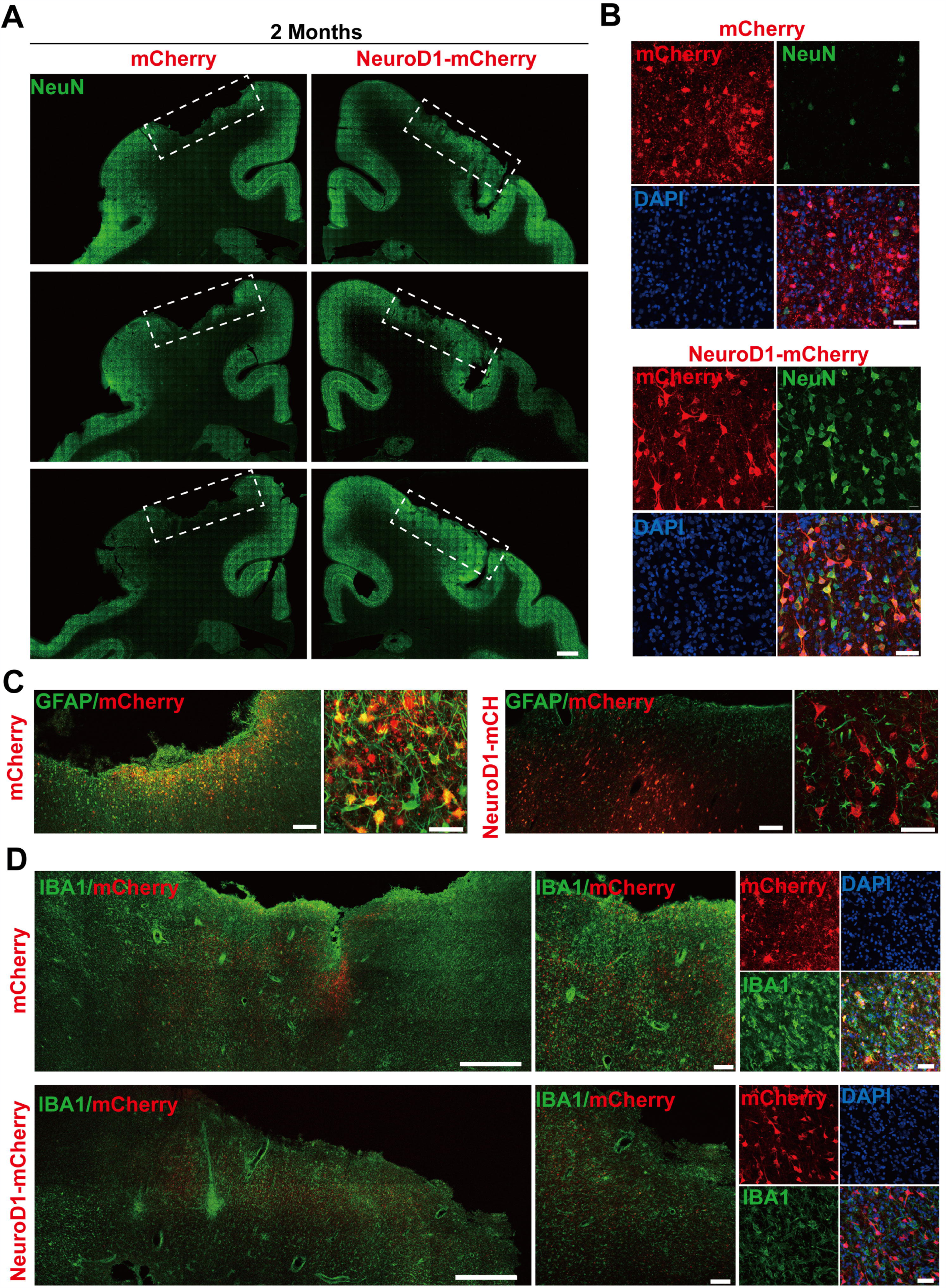
*In vivo* neuroregeneration and reduced inflammation after cell conversion in NHP ischemic stroke model. **(A)** Serial brain sections across the injury core illustrating the neuronal density (NeuN staining) in the monkey cortex following ischemic injury and viral injection. Note that both NeuN signal and cortical tissue were significantly impaired in the mCherry-injected side (left), but significantly rescued in the NeuroD1-mcherry injected side (right). Viral injection was conducted at 21 days post stroke (dps), and immunostaining analysis was performed at 2 months post viral injection (for all panels A to D). Scale bar, 2000 μm. **(B)** Representative high magnification images of neuronal density (NeuN, green) and viral infection (mCherry, red) in the monkey cortex. NeuroD1-mCherry infected areas always showed a significantly increased neuronal density. Nuclei were counterstained with DAPI (blue). Scale bar, 50 μm. **(C)** Both low and high magnification images illustrating the astrocytes (GFAP, green) infected by control virus mCherry alone (red, left panels), but rarely in the NeuroD1-mCherry infected areas (right panels). Note that while rarely co-localizing with NeuroD1-mCherry, GFAP+ astrocytes always persisted in the converted area and even showed less reactive morphology, indicating that astrocytes were not depleted after conversion. Scale bar, 200 µm (low mag), 20 µm (high mag). **(D)** Representative images in low and high magnification illustrate a reduction of reactive microglia (Iba1, green) in the NeuroD1-infected areas (bottom row) following ischemic stroke, comparing to the control side (top row). Nuclei are DAPI stained (blue). Scale bar, 1000 µm (low mag, left panels), 200 µm (higher mag, middle panels), 50 µm (highest mag, right panels).

### *In Vivo* Neuroregeneration after ischemic injury in NHP model

With the successful demonstration of NeuroD1-mediated astrocyte-to-neuron conversion in the monkey cortex, we next investigated whether such *in vivo* neuroregeneration technology can be used to repair damaged brains in NHPs. Since our ultimate goal is to help patients with brain disorders, we decided to pick a neurological condition that is prevalent among patients but still lack effective treatment. Stroke is afflicting millions of patients worldwide but few clinical trials have succeeded in the past (Diener et al., 2008; O’Collins et al., 2006; Percie du Sert et al., 2017; Turner et al., 2013). As the first proof-of-concept study in NHPs to test *in vivo* cell conversion technology, we employed a focal stroke model through intracranial injection of endothelin-1 (ET-1) to induce blood vessel constriction in the motor cortex of Rhesus Macaque monkeys. Although MCAO model is widely used in rodent animals, previous studies in NHPs reported high mortality rate of MCAO model and large variations in the infarct size (Cook and Tymianski, 2012). In contrast, we had established successful ET-1-induced focal stroke model with very low mortality rate and consistent cortical damage in rodents (Chen et al., 2019). In accordance to our rodent studies, when injecting ET-1 into the motor cortex of monkeys, we also observed significant tissue damage at 3 weeks after ischemic injury (Fig. S1A). Specifically, compared to the non-stroke control cortex (Fig. S1A, top row), ET-1 injection caused significant tissue damage as revealed by immunostaining of NeuN, GFAP, and Iba1 at 3 weeks after focal ischemic stroke (Fig. S1A, bottom row). Note that, in ET-1 injected areas, NeuN significantly reduced across all injury areas, whereas GFAP signal also reduced in the lesion core areas but significantly increased in the peri-infarct areas, suggesting that astrocytes became reactive at 3 weeks following ischemic stroke. On the other hand, Iba1 staining showed significant increase in both the lesion core and the peri-infarct areas, suggesting a significant increase of neuroinflammation following ischemic stroke.

After establishing the focal ischemic injury model in the monkey cortex, we examined the effect of NeuroD1-mediated cell conversion on the injured cortical areas. We injected ET-1 (2 μg/μl; 2.5 μl each site; 6 sites for left cortex, with 5 mm apart between two sites, and 6 sites for right cortex) (Fig. S1B, red box) into both left side and right side of the monkey motor cortex to induce focal ischemic stroke. 3 weeks later, control mCherry AAV was injected into one side of the cortex and NeuroD1-mCherry AAV was injected into the other side of the cortex, both into the previous ET-1 injection sites. We then performed a series of immunostaining to investigate the neuronal and glial properties in the NeuroD1-infected cortical tissue versus the control mCherry-infected tissue at 2 month, 4 month, 6 month, and 1 year following a single dose of viral injection (Fig. S1B). Fig. S1C illustrates the gross morphology of monkey cortex after dissection at 2 months following viral injection. The control mCherry-injected site had visible cortical damage, whereas the NeuroD1-injected side showed much better tissue preservation (Fig. S1C, top row). This is confirmed by NeuN immunostaining (Fig. S1C, bottom row), which showed clear tissue loss in the mCherry-injected site but not NeuroD1-injected site. Similar results were also observed at other samples of 4 months and 1 year after viral injection (Fig. S1D-E). Together, ET-1 injection induces severe tissue loss in the monkey cortex, but much milder tissue damage after NeuroD1-treatment.

To investigate the NeuroD1 treatment effect, we performed a series of immunostaining including NeuN, GFAP, and Iba1 to understand the overall neuronal and glial morphology in the stroke areas. Figure 3 illustrates one monkey (monkey ID #07041) at 2 months after viral infection. In the control side injected with mCherry alone (Fig. 3A, serial sections in left column), we found a significant loss of NeuN signal as expected following ischemic injury. However, in the NeuroD1-mCherry injected side, NeuN staining showed much better NeuN signal in a serial sections (Fig. 3A, right column). Enlarged images in Fig. 3B illustrate the significant difference in NeuN signal in the mCherry-infected areas (Fig. 3B, top 4 panels) versus the NeuroD1-infected area (Fig. 3B, bottom 4 panels). Note that most of the NeuroD1-mCherry infected cells were NeuN-positive, suggesting that many converted neurons contributed to the increased neuronal density in the ischemic injured monkey cortex. GFAP immunostaining confirmed that while many mCherry-infected cells remained GFAP+ astrocytes in the control side (Fig. 3C, left panels), the NeuroD1-mCherry infected cells were no longer astrocytes (lost GFAP signal) but showed neuronal morphology (Fig. 3C, right panels). Furthermore, immunostaining with microglia marker Iba1 also revealed that in comparison with the control mCherry-side (Fig. 3D, top row), the NeuroD1-side showed a significant reduction of Iba1 signal (Fig. 3D, bottom row). Similar results were also observed at 4 months (Fig. 4A-E, monkey ID #060107) and 6 months (Fig. 4F-G, monkey ID #04041) following viral infection. Specifically, at 4 months following viral infection, the NeuN signal in the control mCherry side was weaker than that in the NeuroD1-mCherry side (Fig. 4A). The NeuroD1 signal was clearly detected in the NeuroD1-mCherry infected cells (Fig. 4C, green signal) but not in the control side (Fig. 4B). Immunostaining with Iba1 also revealed that compared to the mCherry-infected control areas (Fig. 4D, green signal), there was a significant reduction of reactive microglia in the NeuroD1-mCherry infected areas (Fig. 4E, green signal). At 6 months following viral infection, in comparison to the control mCherry side (Fig. 4F, left panels), we also observed a reduction of Iba1-labeled microglia (green signal) in the NeuroD1-mCherry infected areas (Fig. 4F, right panels). Interestingly, when we investigated parvalbumin-positive (PV) GABAergic interneurons, which are often vulnerable after ischemic injury (Johansen et al., 1990; Povysheva et al., 2019; Tortosa and Ferrer, 1993; Wang, 2003), we found that many PV neurons were surprisingly protected in the vicinity of NeuroD1-mCherry infected areas (Fig. 4G, right panels, comparing to the left control side). Together, these results suggest that NeuroD1-mediated *in vivo* AtN conversion not only generated new neurons in the ischemic injured areas in the monkey cortex, but also reduced microglia and protected GABAergic neurons after ischemic injury.

**Figure 4.**
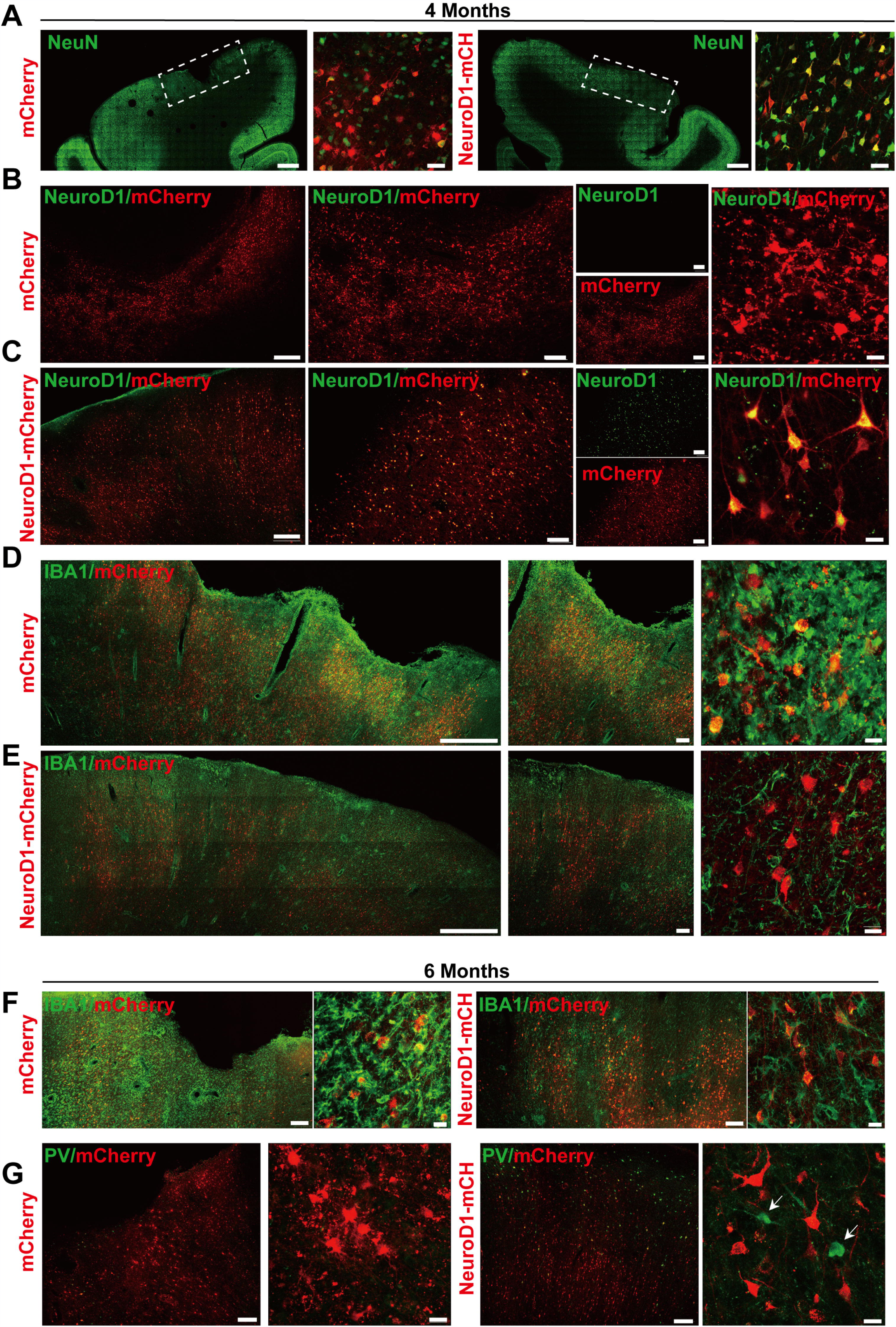
Long-term effect of NeuroD1-treatment in NHP ischemic stroke model. **(A)** Representative images illustrating neuronal density (NeuN, green), cortical tissue integrity, and viral infected cell morphology at 4 months post viral injection (virus injected at 21 dps, for panels A to E). Scale bars, 2000 μm for low mag, 50 μm for high mag. **(B, C)** Identification of NeuroD1 (green) expression at low and high magnification among viral infected cells. As expected, NeroD1 signal (green) was only detected in the NeuroD1-mCherry infected side (C), but not the control side (B). Note that the NeuroD1-mCherry infected cells displayed clear neuronal morphology. Scale bar, 500 µm (low mag, left 4 panels), 200 µm (higher mag, middle 4 panels), 20 µm (highest mag, right 2 panels). **(D, E)** Representative images in low and high magnifications illustrating significant reduction of reactive microglia (IBA1, green) in the NeuroD1-infected areas (E), compared to the control mCherry infected areas (D). Scale bar, 1000 µm (low mag, left panels), 200 µm (higher mag, middle panels), 20 µm (highest mag, right panels). **(F)** Representative images illustrating a reduction of reactive microglia (IBA1, green) in NeuroD1-infected areas (right panels), compared to the control mCherry infected areas (left panels). Virus injected at 21 dps, and immunostaining performed at 6 months post viral injection (for both panel F and G). Scale bar, 200 µm (low mag), 20 µm (high mag). (G) Representative images illustrating the protection of parvalbumin (PV) interneurons (green, arrow) in and surrounding the NeuroD1-infected areas (right panels), compared to the control mCherry-infected areas (left panels). Scale bar, 200 µm (low mag), 20 µm (high mag).

### Increased neuronal density after NeuroD1-treatment

We next performed quantitative analysis on the neuronal density in the NeuroD1-treated versus mCherry control areas after ischemic injury. As shown in Fig. 5A, when compared the NeuroD1-mCherry infected areas versus mCherry alone infected areas, the neuronal density (NeuN+ cells) in NeuroD1-infected areas was always higher than the mCherry-infected areas at 2, 4, and 6 months after viral infection (Fig. 5A, 2^nd^ row NeuN signal). Interestingly, NeuroD1 expression (Fig. 5A, 3^rd^ row) at 6-month time point appeared to be reduced comparing to the 2-month and 4-month time points, which is worth of further investigation in future studies. To obtain a more general assessment of the injured cortical tissue beyond the NeuroD1-infected local regions, we adopted a non-biased quantitative analysis method by sampling the ischemic injured large cortical areas in both control and NeuroD1 sides (illustrated in Fig. 5B). A total of 10 monkeys with ischemic injury on both sides of the motor cortex followed with viral injection (mCherry on one side, and NeuroD1 on the other side) were analyzed using the same method (each monkey analyzed 6 slices on each side, and each slice analyzed 30 – 40 local fields). Non-stroke cortical areas were also quantified in the same manner and yielded a neuronal density of 85/field (single section, each field area 0.1 mm^2^) (Fig. 5C). Among 3 monkeys at 2 months post viral infection, one monkey (07041) showed a significant increase of neuronal density in the NeuroD1-treated side compared to the control side, while the other two monkeys did not show big difference between the two sides (Fig. 5D). All three monkeys at 4-month time point (Fig. 5E) and two monkeys at 6-month time point (Fig. 5F) showed significant increase in the neuronal density after NeuroD1-treatment. After one year of viral infection, one monkey showed significant neuronal increase after NeuroD1-treatment while the other one only showed a mild increase (Fig. 5G). The seemingly less recovery at 2-month time point might suggest that the neuronal recovery takes longer time in non-human primates than in rodents (Chen et al., 2019). Overall, among 10 monkeys treated with NeuroD1-based gene therapy, 7 monkeys showed a significant increase of neuronal density after a single dose of viral injection. Even for the 3 monkeys that did not show overall increase of neuronal density, the NeuroD1-infected local cortical areas typically still showed increase in the neuronal density (see Fig. 5A for the NeuroD1-expressing regions).

**Figure 5.**
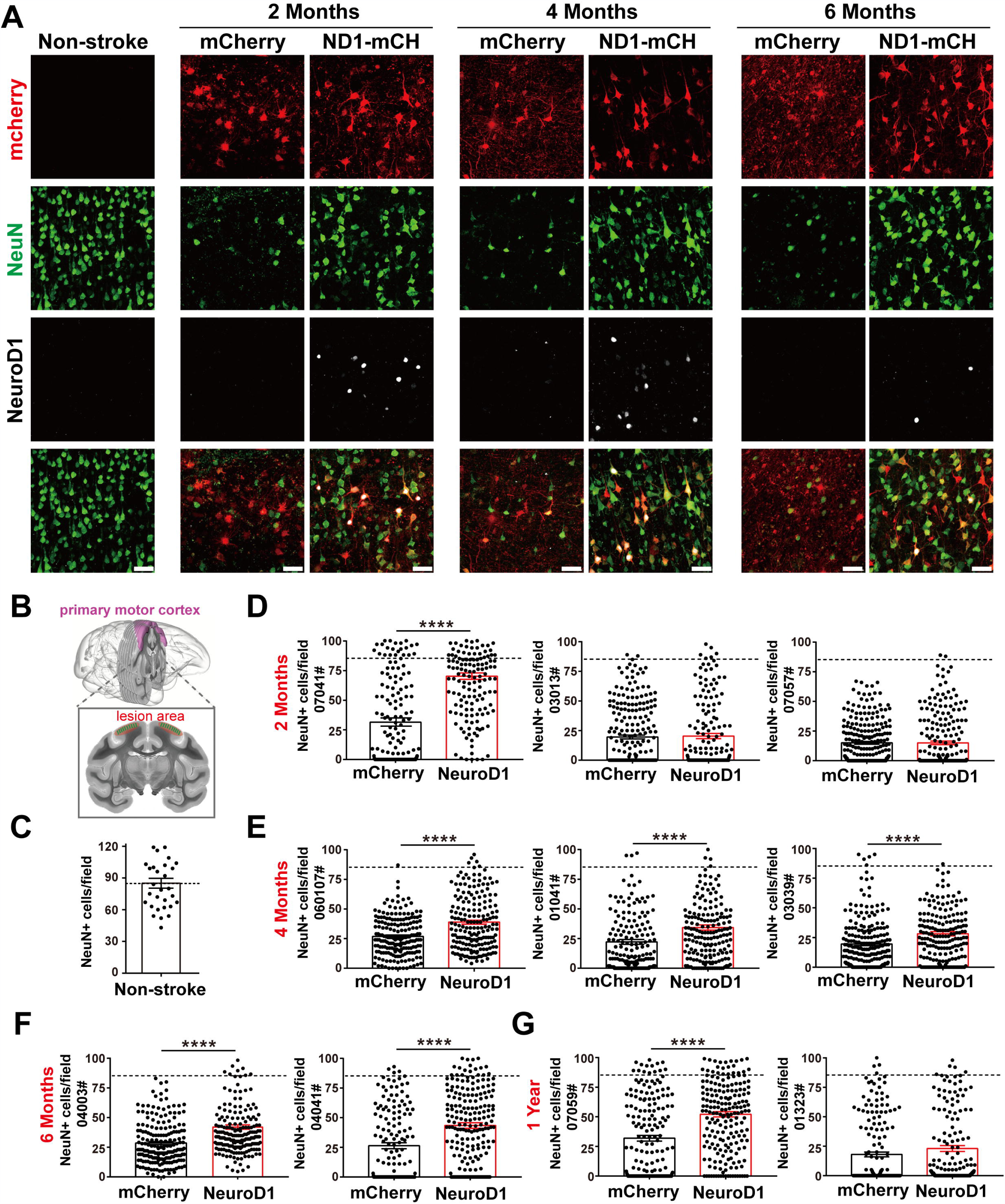
Increased neuronal density after NeuroD1-treatment. **(A)** Representative images showing triple immunostaining of mCherry (red), NeuN (green) and NeuroD1 (white) in non-stroke cortex (left column) and the stroke cortex followed with viral injection (right 6 columns). NeuroD1-infected areas showed a consistent increase in the number of NeuN+ neurons (green) compared to the control side at 2, 4, and 6 months post viral infection. Note that NeuroD1 signal showed a significant decrease at 6 months compared to that at 2 and 4 months post viral infection. Scar bars, 50 μm. **(B)** Schematic illustration of our quantification method. 6 slices across the injury region in the motor cortex of each monkey was picked. 30-40 images of local regions were taken from each side for neuron (NeuN) quantification. **(C)** Quantified data showing the mean number of NeuN+ cells in the motor cortex of non-stroke monkey. Data are represented as mean ± SEM. N = 30 random fields from triplicate slices. Each field = 0.1 mm^2^. **(D-G)** Non-biased quantitative analyses on the neuronal density in the NeuroD1-treated versus control mCherry-treated cortex in 10 monkeys. Viral injection was conducted at 21 days after ischemic injury, and immunostaining was performed at 2, 4, 6 months and 1-year post viral injection. There were 7 out of 10 monkeys showed significantly increased neuronal density after NeuroD1-treatment. Data are represented as mean ± SEM. Each field = 0.1 mm^2^. ****p < .0001 by unpaired two-tailed Student’s *t*-test (for mean value comparison) and Kolmogorov-Smirnov test (for non-parametric ranking order test).

If NeuroD1 converts many reactive astrocytes into neurons, it might raise a question whether such astrocyte-to-neuron conversion would deplete astrocytes in the converted areas. To address this concern, we performed GFAP immunostaining to examine astrocytes in the viral infected areas (Fig. S2). Compared to the ramified resting astrocytes (green signal) in the non-injured areas (Fig. S2A), the astrocytes in the injured areas following control viral injection showed highly hypertrophic morphology with astrocytic processes inter-tangled together (Fig. S2B-D, mCherry columns, green signal). In contrast, in NeuroD1-treated areas, GFAP-labeled astrocytes were always detected in the vicinity of NeuroD1-converted neurons at 2 months, 6 months, or 1 year after viral infection, and less reactive in morphology compared to the control group (Fig. S2B-D, NeuroD1-mCherry columns, green signal). Thus, NeuroD1-mediated high efficiency astrocyte-to-neuron conversion not only generates new neurons but also ameliorates reactive astrocytes without depleting the local astrocytes, consistent with the intrinsic property of astrocytes capable to proliferate.

### Regenerating local cortical neurons after ischemic injury

After the discovery of increased neuronal density following NeuroD1-treatment, we further examined synaptic marker SV2 in the viral infected areas. As shown in Fig. 6A-B, at both 2-month and 6-month time points following viral infection, the SV2 immunostaining showed a significant increase of puncta number in the NeuroD1-infected areas compared to the control mCherry-infected areas, consistent with better neuronal recovery after NeuroD1-treatment.

**Figure 6.**
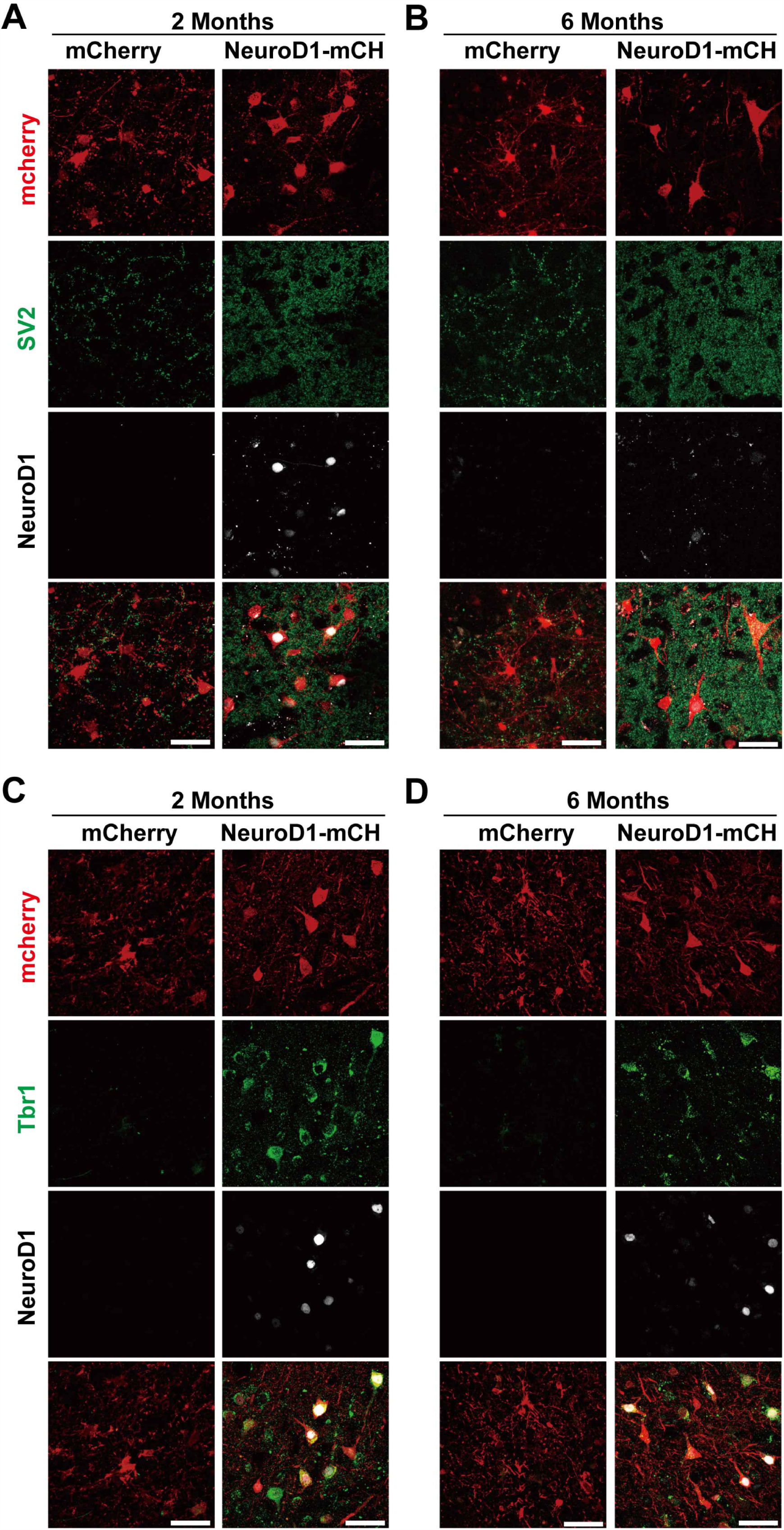
Cortical neuron identity for astrocyte-converted neurons in monkey cortex. **(A-B)** Representative images showing triple immunostaining of mCherry (red), SV2 (green) and NeuroD1 (white) in the NeuroD1-mCherry or mCherry-infected areas at 2 or 6 months post viral injection (21 days after stroke). Data shows significantly increased synaptic puncta (SV2) in the NeuroD1-infected areas, compared to the control mCherry-infected areas. Note a significant decrease of NeuroD1 expression at 6 months following viral infection. Scale bars, 50 μm. **(C-D)** Representative images showing triple immunostaining of mCherry (red), Tbr1 (green) and NeuroD1 (white) in the NeuroD1-mCherry or mCherry-infected areas at 2 or 6 months post viral injection (21 days after stroke). Most of the NeuroD1-expressing neurons were Tbr1+. Scar bars, 50 μm.

We next characterized the neuronal identity of the NeuroD1-converted neurons in the ischemic injured monkey cortex. One difficulty of NHP studies is that many commercially available antibodies that work very well on rodent animals do not work for NHPs. After testing a series of cortical neuron markers, including Tbr1, CTIP2, and CUX2, we found that Tbr1 antibodies (Abcam, catalog #ab31940) showed positive signals in the Rhesus Macaque monkey brains. Importantly, at both 2-month and 6-month time points following viral infection, the NeuroD1-converted neurons were mostly immunopositive for Tbr1 signal (Fig. 6C-D), suggesting that NeuroD1-based gene therapy can regenerate cortical neurons in the monkey cortex after ischemic injury.

We further performed MAP2 immunostaining to assess the neuronal dendrites in the stroke areas after viral infection (Fig. S3). In normal cortex without injury, neuronal dendrites showed strong MAP2 signal (Fig. S3A). After stroke, MAP2 signal was largely lost in the control mCherry-infected areas (Fig. S3B, C, mCherry columns). However, in NeuroD1-infected areas, MAP2 was partially recovered at 2 months and significantly recovered at 6 months post viral infection (Fig. S3B, C, NeuroD1-mCherry columns). In supporting the immunostaining results, we also performed magnetic resonance imaging (MRI) analysis in one monkey at 1 year following ischemic stroke (Fig. S4). Consistent with immunostaining analyses, MRI analysis also revealed tissue loss in the mCherry-infected side (Fig. S4B, yellow arrow) but significant tissue preservation at the NeuroD1-infected side (Fig. S4B, black arrow). Note that the NeuroD1-infected areas did show some kind of tissue abnormality, indicating that it had experienced ischemic injury before. Together, these results suggest that AAV NeuroD1-based gene therapy can regenerate cortical neurons and repair cortical tissue following ischemic injury.

### Protection of GABAergic neurons after astrocyte-to-neuron conversion

As mentioned in Fig. 4, we observed some PV neurons being protected in the NeuroD1-treated group (Fig. 4G). It has been widely reported that ischemic injury tends to severely damage GABAergic neurons (Johansen et al., 1990; Povysheva et al., 2019; Tortosa and Ferrer, 1993; Wang, 2003). In accordance, we also found that compared to non-injured monkey cortex (Fig. 7A, left column), ischemic injury caused a significant loss of PV signal in the mCherry control group (Fig. 7A, mCherry columns, green signal). In contrast, in NeuroD1-treated side, many PV-positive neurons survived in the injured areas (Fig. 7A, ND1-mCH columns, green signal). Interestingly, while the majority of NeuroD1-converted neurons were not PV+ GABAergic neurons, a small number of PV+ neurons in the NeuroD1-infected areas appeared to be mCherry+, consistent with our findings in the rodent animals that NeuroD1 overexpression can generate a small number of GABAergic neurons in the cortex (Guo et al., 2014). Quantitative analysis revealed a significant protection of PV neurons in the NeuroD1-treated side compared to the control mCherry sides. Among the 10 monkeys analyzed from 2 months to 1 year after viral infection, 6 monkeys showed a significant increase in PV+ neurons after NeuroD1-treatment (Fig. 7C-F). The protection of GABAergic neurons after ischemic injury may be critical for preventing further brain damage such as potential epileptic seizures following ischemic stroke (Camilo and Goldstein, 2004; DeFelipe, 1999; Marco et al., 1997; Paz and Huguenard, 2015).

**Figure 7.**
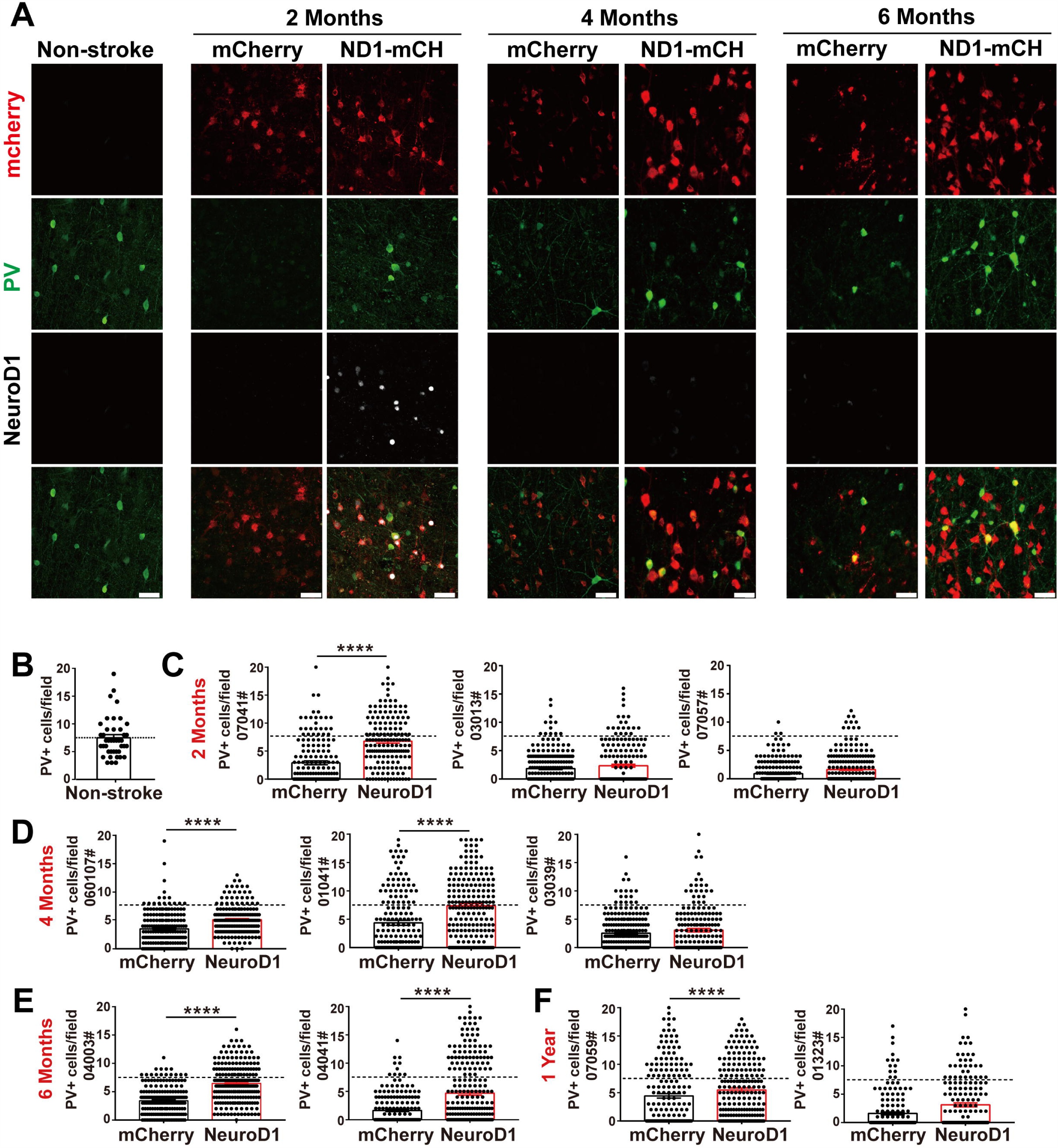
Protection of GABAergic neurons after astrocyte-to-neuron conversion. **(A)** Representative images showing triple immunostaining for mCherry (red), PV (green) and NeuroD1 (white) in non-stroke cortex (left column) and the NeuroD1-mCherry or mCherry-infected areas after stroke. Note a consistent increase of PV+ interneurons in NeuroD1-infected areas compared to the control mCherry-infected areas at 2, 4, and 6 months post viral infection (21 days after stroke). NeuroD1 expression decreased at 4 and 6 months after infection, but mCherry signal was still strong in the viral infected cells. Scar bars, 50 μm. **(B)** Quantified data showing the mean number of PV+ cells in the motor cortex of non-stroke monkey. Data are represented as mean ± SEM. N = 30 random fields from triplicate slices. Each field = 0.1 mm^2^. **(C)** Quantitative analysis on the PV+ neuronal density in the NeuroD1-treated versus control mCherry-treated cortex in 10 monkeys. There were 6 out of 10 monkeys showed significant increase of PV+ neurons after NeuroD1-treatment. Viral injection at 21 days following ischemic injury, and immunostaining at 2, 4, 6 months and 1-year post viral injection. 6 slices on each side were quantified from each monkey, and 30-40 local regions from each slice were quantified. Data are represented as mean ± SEM. ****p < .0001 by unpaired two-tailed Student’s t test (for mean value comparison) and Kolmogorov-Smirnov test (for all dotted points).

### Broad time windows of NeuroD1-treatment after ischemic injury

Finally, we injected NeuroD1 AAV at different time windows following ischemic injury. Besides viral injection at 21 days after stroke in 10 monkeys shown above, we also injected mCherry control or NeuroD1-mCherry AAV at 10 days post ischemic injury (10 dpi) (Fig. 8 and Fig. S5) or 30 days post ischemic injury (Fig. S6 and Fig. S7). Fig. 8A illustrates the experimental design on two monkeys injected AAV at 10 days post stroke. Similar to the experiments shown above for viral injection at 21 days post stroke, we consistently observed increased neuronal density (Fig. 8B, NeuN staining) and better recovery of MAP2 signal in the NeuroD1-infected areas (Fig. 8C, green signal) at both 2-month and 4-month time points following viral infection. Furthermore, the NeuroD1-infected areas also showed less reactive microglia (Fig. S5A) and more preserved PV-positive neurons (Fig. S5B). The same results were also observed in two monkeys injected AAV at 30 days following ischemic stroke (Fig. S6A). Specifically, NeuroD1-treatment showed significant increase in neuronal density (Fig. S6B, NeuN signal in green), better recovery in MAP2-labeled dendrites (Fig. S6C, green signal), less reactive microglia (Fig. S7A), and more preserved PV-positive neurons (Fig. S7B). While more experiments are necessary to test the optimal time window, our results suggest that NeuroD1 AAV-based gene therapy may have a broad treatment window ranging from 10 to 30 days post ischemic stroke to have effective therapeutic intervention.

**Figure 8.**
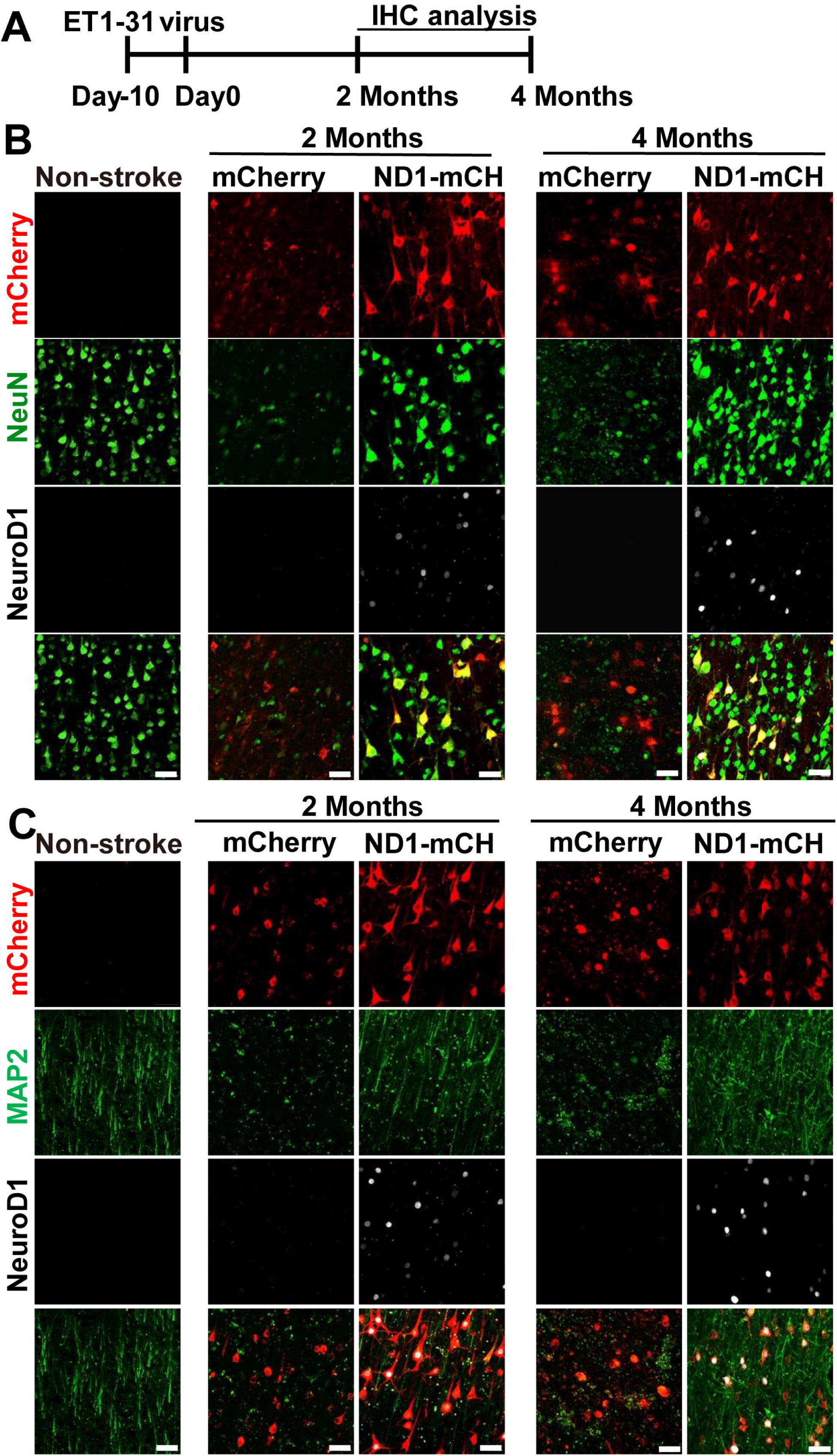
Beneficial effects of NeuroD1-based gene therapy delivered at 10 days after ischemic stroke in monkey cortex. **(A)** Experimental design for NeuroD1 treatment at 10 days after ischemic stroke. **(B)** Representative images showing mCherry (red), NeuN (green) and NeuroD1 (white) expression pattern in non-stroke cortex (left column) or in ischemic cortex after virus infection (right 4 columns). NeuroD1-infected areas consistently showed higher neuronal density. Scar bars, 50 μm. **(C)** Representative images showing mCherry (red), MAP2 (green), and NeuroD1 (white) expression pattern in non-stroke cortex (left column) or in ischemic cortex after viral injection (right 4 columns). The neuronal dendritic marker MAP2 signal increased significantly in NeuroD1-infected areas, compared to the control mCherry-infected areas. Scar bars, 50 μm.

## DISCUSSION

In this study, we demonstrate that ectopic expression of a single neural transcription factor NeuroD1 in the reactive astrocytes of monkey cortex after stroke can successfully convert astrocytes into neurons. This is built upon our earlier success in mouse and rat brains *in vivo* as well as human astrocytes *in vitro* that NeuroD1 can mediate powerful AtN conversion to regenerate a large number of functional new neurons for brain repair (Chen et al., 2019; Guo et al., 2014; Zhang et al., 2018). A successful application of our NeuroD1 AAV-based gene therapy in a NHP model bridges an important gap toward future clinical trials in human patients.

### *In vivo* cell conversion in rodent versus NHP models

Neuronal loss is the most common cause of brain functional deficits after acute injury such as stroke and concussion as well as chronic neurodegenerative disorders such as Alzheimer’s and Parkinson’s disease. Neuroregeneration in adult mammalian CNS has been proved to be one of the most difficult tasks in the entire regenerative medicine field, largely because neurons cannot divide to regenerate themselves and external cell transplantation yields very low number of functional new neurons (Goldman, 2016). To overcome the limitations of cell transplantation therapy, we, together with other research groups, have developed *in vivo* cell conversion technology to regenerate functional new neurons from endogenous glial cells for brain repair (Chen et al., 2019; Gascon et al., 2016; Guo et al., 2014; Karow et al., 2018; Liu et al., 2015; Niu et al., 2015; Niu et al., 2018; Pereira et al., 2017; Torper et al., 2015; Wang et al., 2016; Zhang et al., 2018). While stab injury model has been commonly used in various *in vivo* cell conversion studies, we have previously demonstrated in a mouse Alzheimer’s disease model that NeuroD1 can convert reactive astrocytes into functional neurons in 14-month old AD mouse brains (Guo et al., 2014). Recently, we have further demonstrated in a mouse stroke model that NeuroD1-based gene therapy can successfully convert reactive astrocytes into functional neurons and promote functional recovery (Chen et al., 2019). Other group also reported that striatal astrocytes can be converted into dopaminergic neurons and promote behavioral recovery in a mouse PD model (Rivetti di Val Cervo et al., 2017). These successful *in vivo* cell conversion studies in various rodent animal models suggest a great potential in neuroregeneration and neural repair. Nevertheless, since so many clinical trials on CNS disorders have failed in the past based on successful preclinical studies in rodent animal data, we believe that it is essential to establish solid foundation in NHP models before starting clinical translation for human brain repair. Some successful discoveries made in the rodent animal models may not be applicable to primates. For example, a recent study generated a new serotype of AAV PHP.eb that showed remarkable permeability through blood-brain-barrier in mice and could infect essentially the entire brain with high efficiency (Chan et al., 2017). However, it was later found that such AAV serotype was better suited for rodent animals only, and did not show high infection efficiency in NHPs (Liguore et al., 2019), suggesting that it would be a disaster should someone jump from mouse success directly to human clinical trials using this viral serotype. For stroke studies, it is also known that the blood vessel branching network and lateral circulation in rodents are very different from primates (Cuccione et al., 2016; Du et al., 2016; Masakatsu et al., 2003; Shinichi et al., 2004; Shunichi and Zoppo, 2003). Therefore, the ischemic injury and its post-injury compensation may well be different between rodents and primates. Our current study extends our previous success on *in vivo* AtN conversion achieved in rodent animals (Chen et al., 2019; Guo et al., 2014; Zhang et al., 2018), and provides direct evidence that NeuroD1-based gene therapy can convert reactive astrocytes into neurons in monkey brains suffered from ischemic injury. This *in vivo* NHP model fills the gap between *in vivo* rodent model and *in vitro* human glial cell culture model with regard to AtN conversion, making one leap toward future clinical trials. In deed, current gene therapy products on the market including Luxturna from Spark Therapeutics and Avexis ZOLGENSMA have both obtained successful NHP data (Jacobson et al., 2006; Passini et al., 2014; Towne et al., 2010) before conducting human clinical trials.

One unexpected observation in our NHP study on NeuroD1-mediated AtN conversion is the gradual decrease of NeuroD1 expression level at 6 months after viral infection, which was not observed in mice, highlighting the difference between rodents and primates. As a neural transcription factor, NeuroD1 is known to be expressed in a subset of neural stem cells during early brain development and then decrease in the adult brain (Cho and Tsai, 2004; Gaudillière et al., 2004; Lee et al., 2010; MuñOz et al., 2010; Pataskar et al., 2016; Pleasure et al., 2000), except the adult hippocampus in mice (Gao et al., 2009; Tomoko et al., 2009). In both mouse and monkey brains, immunostaining with NeuroD1 antibodies always detected very low level of NeuroD1 signal in the cortical neurons. In NeuroD1-infected cells, NeuroD1 signal is typically very high compared to the non-infected cells. This is true in monkey brains at 2-month and 4-month time points after viral infection, but we observed a significant decrease in NeuroD1 signal within many NeuroD1-mCherry infected cells at 6-month time point. While the precise mechanism of such decrease of NeuroD1 expression after long-term neuronal conversion and maturation requires further investigation, it suggests that the NeuroD1 expression from the AAV episome may be significantly downregulated in mature neurons, for example through phosphorylation regulated silencing mechanism. Such spontaneous downregulation of NeuroD1 after neuronal conversion might be an advantage for future clinical trials, because NeuroD1 may not stay at a high level in converted neurons.

It is worth to emphasize that there are also many similarities between rodent and NHP models regarding AtN conversion. One of the most consistent findings is that ectopic expression of NeuroD1 in cortical reactive astrocytes after ischemic injury can efficiently convert astrocytes into neurons (90% conversion efficiency) regardless of rodents or NHPs. Another interesting finding is that, while white matter occupies much larger area in monkey cortex, we rarely detect any converted neurons in the monkey white matter, which is consistent with our recent findings in rodent white matter (Liu et al., 2020).

### Broad time window for NeuroD1 AAV-based gene therapy

Previously, there have been a number of stroke studies in NHP models but they are mainly testing drug effects in acute stroke model typically within a few hours after stroke (Cook et al., 2017; Cook et al., 2012; Takamatsu et al., 2001; Yasuhisa et al., 2003). In our current study, we have tested NeuroD1-mediated neuronal conversion at 10, 21, and 30 days after ischemic injury and found that neurons can be regenerated at least one month following stroke, indicating that our *in vivo* cell conversion technology may have a broad time window for stroke treatment. Furthermore, the neuronal recovery after NeuroD1 treatment is long lasting, ranging from 2 months to one year after a single dose of AAV treatment. Such structural improvement at the cellular level may be critical for further functional improvement, which will be tested in ongoing studies.

Interestingly, similar to our recent observations made in mouse stroke studies (Chen et al., 2019), we also observed significant protection of interneurons after NeuroD1 treatment in monkey stroke model. It is known that interneurons are vulnerable under ischemic injury (Johansen et al., 1990; Povysheva et al., 2019; Tortosa and Ferrer, 1993; Wang, 2003). The neuroprotective effect of *in vivo* cell conversion mediated by NeuroD1 may have important implications for therapeutic treatment. The precise mechanisms are to be investigated in further studies, but our observation of reduced reactive microglia at the NeuroD1-infected areas may at least contribute to the protection of interneurons as well as other neurons.

It is worth to mention that one big challenge about working on non-human primates is that many antibodies that work very well for rodent animals often do not work for monkeys. To identify the variety of neuronal subtypes in monkey brains, it is necessary to apply new technologies such as single cell sequencing technology to better characterize the identity of newly converted neurons.

### NeuroD1 AAV as a potential therapeutic agent

Previous clinical trials on stroke repair have largely failed because of lack of direct translation from successful rodent studies into human patients (Turner et al., 2013). A successful demonstration of neural repair in adult NHP models after stroke will be an important step toward potential translation into clinical therapies. Our current study in NHP stroke model, together with our recent rodent stroke study (Chen et al., 2019), shows a consistent pattern that NeuroD1-based gene therapy can regenerate a large number of new neurons in the adult mammalian brains, which is the fundamental first step toward brain repair. The limitation of this current NHP study is that the neuronal recovery at the cellular level needs to be further correlated with behavioral improvement, which requires a large number of NHPs for statistical analysis. Another limitation is that the current stroke model is a focal stroke model, with limited total infarct volume caused by stroke. Further studies will test more severe stroke models such as cerebral artery occlusion or hemorrhagic stroke models in NHPs to better mimic patient conditions in order to develop more suitable therapeutic interventions for patients.

## Materials and methods

### Animal preparation

Eighteen adult rhesus macaques (*Macaca mulatta*), aged from 9 to 21 years old (mean body weight = 8.5 kg), were employed in this study. The monkeys were provided by the Kunming Primate Research Center, Kunming Institute of Zoology, Chinese Academy of Sciences, China. All rhesus macaques were male and healthy without history of neurological diseases, and were housed in individual cages with a standardized light/dark cycle and were fed under standard procedures. All experimental procedures and animal care were approved by the Ethics Committee of Kunming Institute of Zoology and the Kunming Primate Research Center (approval NO. IACUC19007), Chinese Academy of Sciences, performed in accordance with National Institutes of Health guidelines for the care and use of laboratory animals. The animal care facility was accredited by the Association for Assessment and Accreditation of Laboratory Animal Care (AAALAC). One monkey (ID #04013) was used in the preliminary study for the establishment of focal ischemic stroke model, and another monkey (ID #04339) was tested for *in vivo* astrocyte-to-neuron conversion without stroke. Other sixteen monkeys were used for focal ischemic stroke followed by viral injection. To use minimal numbers of monkeys, one side of the monkey cortex was used for control viral injection, and the other side of monkey cortex was injected with NeuroD1 AAV for cell conversion test. Information for these monkeys was listed in Table1.

**Table 1.** Animal summary

| Table 1. Animal summary |  |  |  |  |  |  |
| --- | --- | --- | --- | --- | --- | --- |
| Animals | Sex | Age | Bodyweight(KG) | ET1 injury | Virus | Virus Duration |
| 04339# | M | 12 | 6.7 | —— | AAV-GFAP::Cre+AAV-Flex-CAG-ND1/mCherry+AAV-GFAP::GFP; GFAP::GFP; GFAP::ND1-GFP | 4-6 weeks |
| 04013# | M | 12 | 8.6 | 3 weeks | —— | —— |
| 07057# | M | 9 | 12.0 | 3 weeks | AAV-GFAP::Cre+AAV-Flex-CAG-ND1/mCherry | 2 Mon |
| 07041# | M | 9 | 10.4 | 3 weeks | AAV-GFAP::Cre+AAV-Flex-CAG-ND1/mCherry | 2 Mon |
| 03013# | M | 10 | 12.5 | 3 weeks | AAV-GFAP::Cre+AAV-Flex-CAG-ND1/mCherry | 2 Mon |
| 01041# | M | 15 | 6.7 | 3 weeks | AAV-GFAP::Cre+AAV-Flex-CAG-ND1/mCherry | 4 Mon |
| 060107# | M | 10 | 7.1 | 3 weeks | AAV-GFAP::Cre+AAV-Flex-CAG-ND1/mCherry | 4 Mon |
| 03039# | M | 13 | 11.0 | 3 weeks | AAV-GFAP::Cre+AAV-Flex-CAG-ND1/mCherry | 4 Mon |
| 04003# | M | 12 | 9.6 | 3 weeks | AAV-GFAP::Cre+AAV-Flex-CAG-ND1/mCherry | 6 Mon |
| 04041# | M | 12 | 10.1 | 3 weeks | AAV-GFAP::Cre+AAV-Flex-CAG-ND1/mCherry | 6 Mon |
| 07059# | M | 9 | 8.5 | 3 weeks | AAV-GFAP::Cre+AAV-Flex-CAG-ND1/mCherry | 1 year |
| 01323# | M | 15 | 16.0 | 3 weeks | AAV-GFAP::Cre+AAV-Flex-CAG-ND1/mCherry | 1 year |
| 03021# | M | 14 | 7.1 | 10 days | AAV-GFAP::Cre+AAV-Flex-CAG-ND1/mCherry | 2 Mon |
| 01073# | M | 16 | 6.9 | 10 days | AAV-GFAP::Cre+AAV-Flex-CAG-ND1/mCherry | 2 Mon |
| 05025# | M | 12 | 8.7 | 10 days | AAV-GFAP::Cre+AAV-Flex-CAG-ND1/mCherry | 4 Mon |
| 07089# | M | 10 | 7.7 | 30 days | AAV-GFAP::Cre+AAV-Flex-CAG-ND1/mCherry | 2 Mon |
| 04307# | M | 13 | 7.1 | 30 days | AAV-GFAP::Cre+AAV-Flex-CAG-ND1/mCherry | 2 Mon |
| 96079# | M | 21 | 8.0 | 30 days | AAV-GFAP::Cre+AAV-Flex-CAG-ND1/mCherry | 4 Mon |

### Stroke Model

Food, but not water, was deprived in the evening prior to the surgery day to avoid potential choke of food during anesthesia. Anesthesia was induced with hydrochloric acidulated ketamine (10 mg/kg, intramuscular injection) and maintained with sodium pentobarbital (Merck; 20 mg/kg, i.m.). Atropine (0.02 mg/kg, i.m) was given preoperatively to decrease secretions. Heart rate, blood pressure and blood oxygen saturation were monitored. For cortical injection, the monkey was placed on a warm heated surgical table and mounted on a stereotaxic apparatus (Shenzhen Reward Life Science, Shenzhen, China). The focal ischemic lesion was created in the bilateral primary motor cortex (M1) area. The animals were operated with a midline scalp incision followed by drilling holes on the skulls above motor cortex. Micro-injection of Endothelin-1 (1-31) (Human) (Peptide, 4360-s, Japan) was made at six sites in each side of the M1 area to induce focal ischemic injury according to a combined MRI and Histology Atlas of the Rhesus Monkey Brain in Stereotaxic Coordinates (2nd Edition, Kadharbatcha S. Saleem, Nikos K. Logothetis, 2012) at the following coordinates: (1) anteroposterior (AP), 18.8 mm; mediolateral (ML), 11.2 mm, (2) AP, 17.3 mm; ML, 13.5 mm, (3) AP, 15.7 mm; ML, 16.1 mm, (4) AP, 16.6 mm; ML, 8.9 mm, (5) AP, 15.6 mm; ML, 11.1 mm, (6) AP, 14.4 mm; ML, 13.9 mm. A volume of 2.5 μL (2 μg/μL) of ET-1 was injected at each site using a 10-μL Hamilton syringe and infusion pump (UMP3-2, Micro4, WPI Apparatus, Sarasota, USA) at a speed of 100 nl per minute. Six injections were given 4 mm under the dura of monkey. Needle was inserted 4 mm under the dura and left in place for 5 minutes to allow the needle completely fit into brain tissues. Then the needle was withdrawn to 2 mm under the dura, and started to inject after 2-minute pause. ET-1 was injected at each site in five boluses of 500 nL, allowing a 2-minute pause and 500 μm distance between each bolus. At the end of ET-1 delivery at each site, the needle was left in place for 5 minutes to allow ET-1 solution diffuse away from the needle tip before withdrawal. Then the operative field was washed with saline and the incision was sutured. After operation, the monkeys received antibiotic treatment of penicillin sodium (100,000 U/kg, intramuscular injection daily) for 3 days to prevent infection.

### Plasmid construction and AAV virus production

To produce adeno-associated virus (AAV) vectors, ND1-P2A-mCherry was inserted into pAAV-FLEX-GFP vector (Addgene) to replace GFP to generate pAAV-Flex-CAG-ND1-P2A-mCherry. mCherry was inserted into pAAV-FLEX-GFP to replace GFP and was used as a negative control. Human GFAP promoter (1.6 kb) was inserted into pAAV-MCS (Cell Biolabs) to replace CMV promoter, then Cre was subcloned into pAAV-MCS to generate pAAV-hGFAP-Cre. The backbone of pAAV-GFAP::ND1-GFP and pAAV-GFAP::GFP vectors is pAAV-MCS (Cell Biolabs, Inc). CMV promoter was changed to GFAP promoter, and GFP or NeuroD1-GFP was inserted to MCS. Recombinant AAV9 was produced by HEK293T cell line. The purification method used Iodixanol Gradient Solutions (D1556, Optiprep, Sigma) and desalting and concentrating on Merck Amicon Ultra-15 Centrifugal Filters (UFC910008, Millipore). Purified AAV viruses were titered using QuickTiter™ AAV Quantitation Kit (VPK-145, Cell biolabs).

### AAV virus stereotactic injection

Animals were anesthetized as mentioned above before fastened to the stereotaxic instrument. Total 10 μl viruses per site were injected using the Hamilton syringe and infusion pump at a speed of 400 nl per minute. AAV viruses were injected in the same sites of ET-1 injection. To test conversion, AAV viruses were directly injected in non-stroke monkey cortex (monkey ID #04339). Virus stocks were diluted to 10^12^ GC/ml for injection. After injection, the needle was kept in place for five additional minutes and then slowly withdrawn. Then, the wound was cleaned thoroughly and sutured. After surgery, monkeys were given penicillin daily for 3 days to prevent infection.

### MRI Scanning

All magnetic resonance imaging (MRI) scans were performed using the LianYing uMR770 3.0-T scanner. Two monkeys were arranged for MRI analysis one year after virus injection. Prior to the MRI scan, monkeys were anesthetized intramuscularly as previously described. T2-weighted imaging was acquired in the axial plane and included the following sequences: repetition time (TR) = 2300.0 ms; echo time (TE) = 382.7 ms; field of view (FOV) = 120 × 120 mm (T1), 240 × 240 mm (T2); Matrix = 240 × 240 (T1), 320 × 320 (T2); slice thickness 0.5 mm, no gap.

### Immunohistochemistry

For brain section staining, the monkeys were deeply anesthetized with sodium pentobarbital and perfused transcardially with phosphate-buffered saline (PBS, pH 7.4) followed by 4% (W/V) paraformaldehyde (PFA) in phosphate-buffered saline to fix the brain. Brains were carefully removed from the skull and post fixed in 4% PFA for further 3 days at 4 °C, and then cut in 40 µm thickness by a vibratome (VT-1000S; Leica).

For immunofluorescence, coronal brain sections were first pretreated in 0.3% Triton X-100 in PBS for 30 min, followed by incubation in 5% bovine serum albumin (BSA) and 0.1% Triton X-100 in PBS for 1 hour at room temperature. For double or triple immunofluorescence, primary antibodies were simultaneously incubated overnight at 4 °C in 1% BSA and 0.1% Triton X-100 in PBS. Primary antibodies were used as follows: mouse anti-GFAP (1:1000, SMI-21R, Biolegend), goat anti-NeuroD1 (1:1000, sc-1084, SantaCruz), rabbit anti-NeuN (1:1000, ABN78, Millipore), mouse anti-NeuN (1:500, MAB377, Millipore), mouse anti-MAP2 (1:500, M4403, Sigma), chicken anti-GFAP (1:1000, AB5541 Millipore), rabbit anti-Iba1 (1:1000, 019-19741, Wako), rabbit anti-AQP4 (1:500, AB3594, Millipore), rabbit anti-Parvalbumin (1:1000, ab11427, Abcam), rabbit anti-GFP (1:1000, sc-8334, SantaCruz), chicken anti-GFP (1:1000, ab13970, Abcam), rabbit anti-mCherry (1:1000, ab167453, Abcam), chicken anti-mCherry (1:500, 205402, Abcam), mouse anti-SV2 (1:1000, SV2, DSHB), rabbit anti-Tbr1 (1:500, ad31940, abcam). The next day, primary antibodies were washed off and proper fluorophore-conjugated secondary antibodies were used, Cy2-, Cy3-, and Cy5-conjugated secondary antibodies were obtained from Jackson ImmunoResearch (1:500). Alexa Fluor 488-, Alexa Fluor 594-, and Alexa Fluor 647-conjugated secondary antibodies were obtained from Thermo Fisher Scientific (1:1000). After additional washing in PBS, sections were counterstained with DAPI (4,6-diamidino-2-phenylindole, D8200, Solarbio) in PBS for 10 min. The sections were then rinsed 3 times with PBS and mounted onto a glass slide with a Fluorescent Mounting Media (71-00-16, KPL). Stained sections were examined and photographed with Nikon A1+ or Zeiss LSM880 confocal microscopy. Z-stacks of digital images were acquired using Nikon A1+ or Zeiss LSM880 confocal microscopy.

For DAB staining, endogenous peroxidase activity was quenched through incubation of sections in 3% H2O2 for 10 minutes at room temperature prior to washing with PBS, and sequential incubations with blocking serum and primary antibodies (overnight 4 °C) were performed. Secondary peroxidase-labeled affinity purified goat anti-mouse (474-1806, KPL) and goat anti-rabbit antibodies (474-1516, KPL) were used at 1/500 dilution for 2 hours. After washing in PBS, the sections were developed with diaminobenzidine (DAB). The sections were rinsed with PBS and then mounted on slides, dehydrated, and coversliped. Sections were viewed with microscope (Olympus, CX41; camera: Olympus DP25; software: CellSens Entry 1.4.1; Japan).

### Statistical analyses

All images for quantitative analyses were acquired by a confocal microscope (Zeiss LSM880, Jena, Germany). Data was presented as mean ± SEM. Unpaired Student’s *t*-Test was used (Graphpad Prism 7, GraphPad Software Inc., San Diego, CA, USA) to compare datasets from experimental and control groups.

## ACKNOWLEDGMENTS

This work was supported by Yunnan Major Project of Science and Technology (2014FC005), Yunnan Key Program of Science and Technology (2017FA042), and National Science Foundation of China (31571109). G.C. is a co-founder of NeuExcell Therapeutics Inc.

## AUTHOR CONTRIBUTION

G.C. conceived and supervised the entire project, analyzed the data, and wrote the manuscript. LJG performed the majority of the experiments with significant help from FHY, analyzed the data, and participated in writing the manuscript. LJG and FHY performed monkey brain surgery experiments with the assistance of NHC and JF. LJG made viruses and performed the MRI experiment. LJG, FHY, MJ and JF all contributed to immunostaining experiment. XTH and JHW participated in the discussion of the experiments.

