## supplemental data for "*In Vivo* Neuroregeneration to Treat Ischemic Stroke in Adult Non-Human Primate Brains through NeuroD1 AAV-based Gene Therapy"

### Supplementary Figures and Legends:

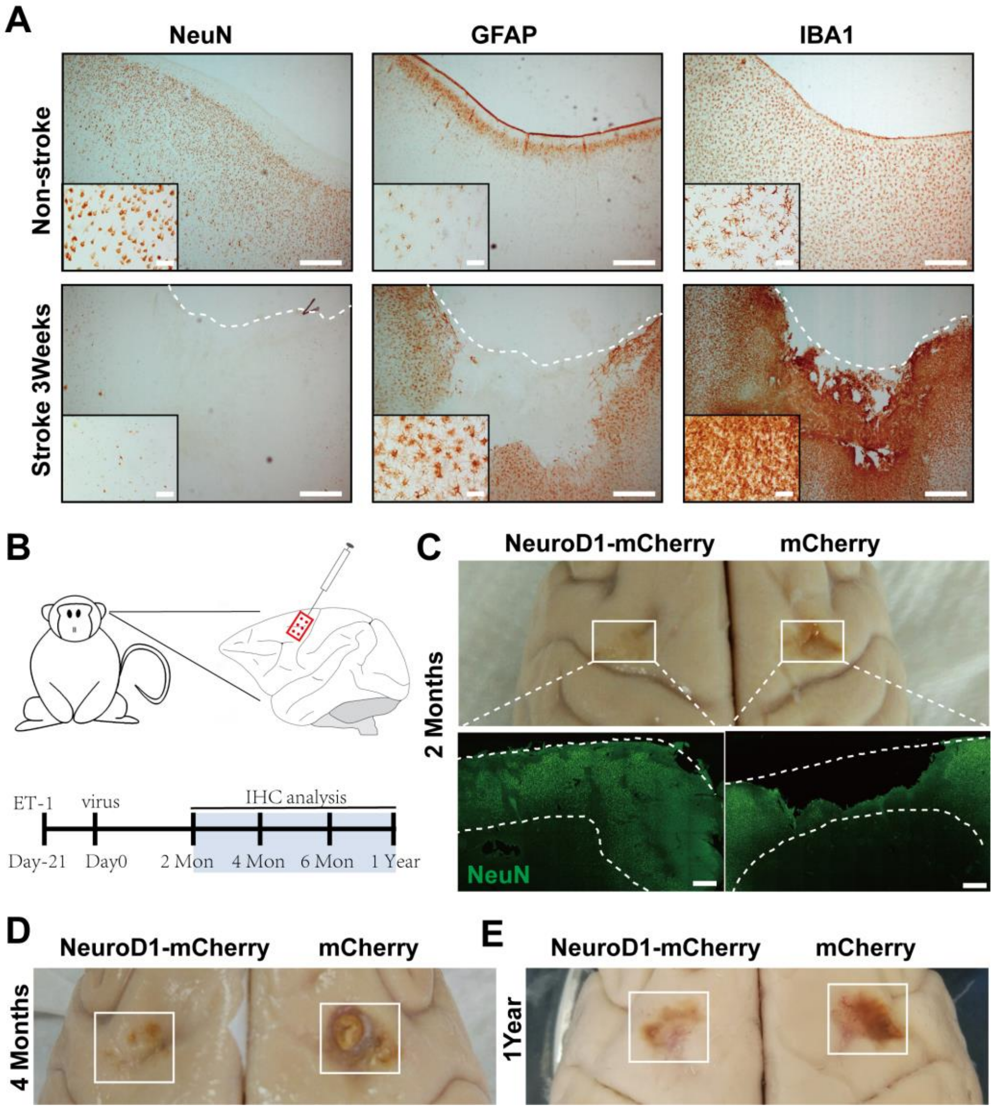


**Figure S1. Establishment of NHP ischemic cortical stroke model and brain repair by NeuroD1 AAV-based gene therapy.**
**(A)** Representative images showing NeuN, GFAP and Iba1 immunostaining from cortical sections of non-stroke cortex (top row) and the endothelin-1 (ET-1, 1-31)

induced ischemic stroke cortex (bottom row). Note that loss of neurons and gliosis occurred at 3 weeks after focal ischemic injury. Scale bars, 500 μm (low mag), 50 μm (high mag).
**(B)** Schematic diagram illustrating experimental design for the majority of NeuroD1 gene therapy intervention.
**(C)** Representative images showing brain tissue integrity at 2 months following viral injection after stroke. Note a significant tissue damage in the mCherry-injected side (top right) compared to the NeuroD1-injected side (top left), which was further confirmed by immunostaining of NeuN (green) (bottom row). Dashed lines indicate cortical areas. Scale bars, 1000 μm.
**(D-E)** Representative images illustrate less tissue damage in NeuroD1-treated side compared to control mCherry-treated side at 4 months (D) and 1 year (E) following viral injection after stroke.

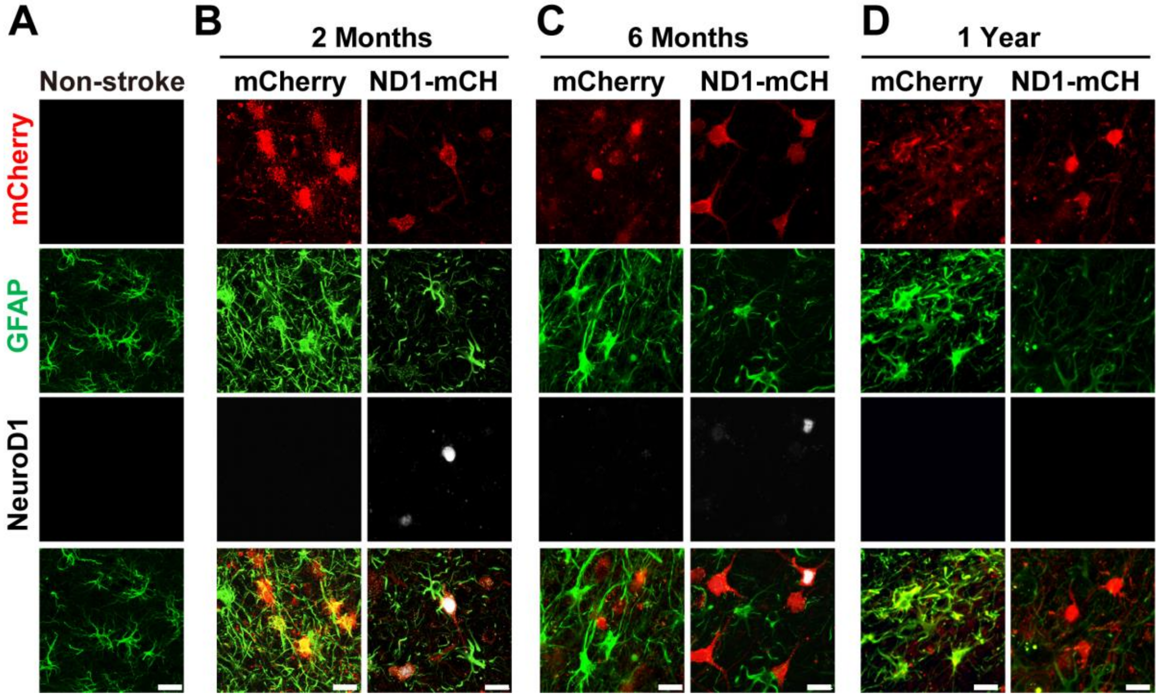


**Figure S2. Astrocytes are not depleted after NeuroD1-mediated conversion. (A-D)** Representative images showing triple immunostaining of mCherry (red), GFAP (green), and NeuroD1 (white) in brain sections obtained from non-stroke monkey (A) and stroke monkeys at 2 months (B), 6months (C), and 1 year (D) after viral infection. Note that in NeuroD1-infected areas, astrocytes were always present and their morphology was less reactive, accompanied with less GFAP protein level compared to the control mCherry-infected cortex. Scar bars, 20 μm.

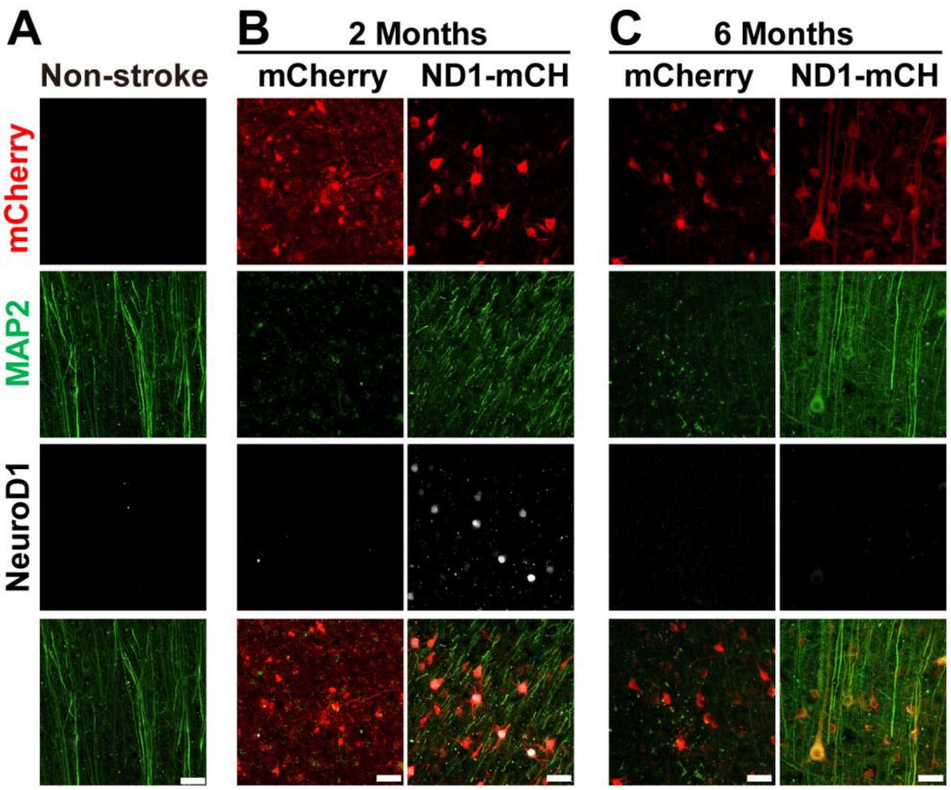


**Figure S3. NeuroD1-treatment rescued neuronal dendritic morphology after ischemic stroke in monkey cortex.**

**(A-C)** Representative images showing triple immunostaining of mCherry (red), MAP2 (green) and NeuroD1 (white) in brain sections obtained from non-stroke cortex (A) and ischemic injured cortex at 2 months (B) and 6 months (C) after viral injection. Note that NeuroD1 expression at 6 months after viral infection was significantly reduced, suggesting a potential self-downregulation mechanism. Neuronal dendrites labeled by MAP2 were significantly rescued in the NeuroD1 group compared to the control group. Scar bars, 50 μm.

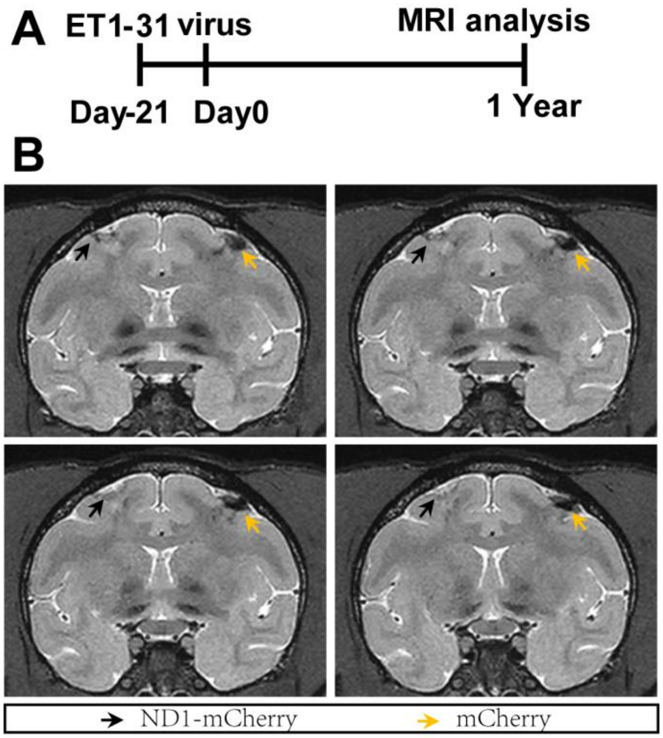


**Figure S4. Brain tissue repair of monkey cortex by NeuroD1-treatment revealed by T2-weighted magnetic resonance imaging (MRI) after ischemic injury.**
**(A)** Timeline for experimental design.
**(B)** T2-weighted MRI images of a monkey brain showing ischemic injured motor cortex at 1 year after AAV viral injection in the control side (right， yellow arrow) and NeuroD1 treatment side (left， black arrow). Arrow points to the location of lesion.

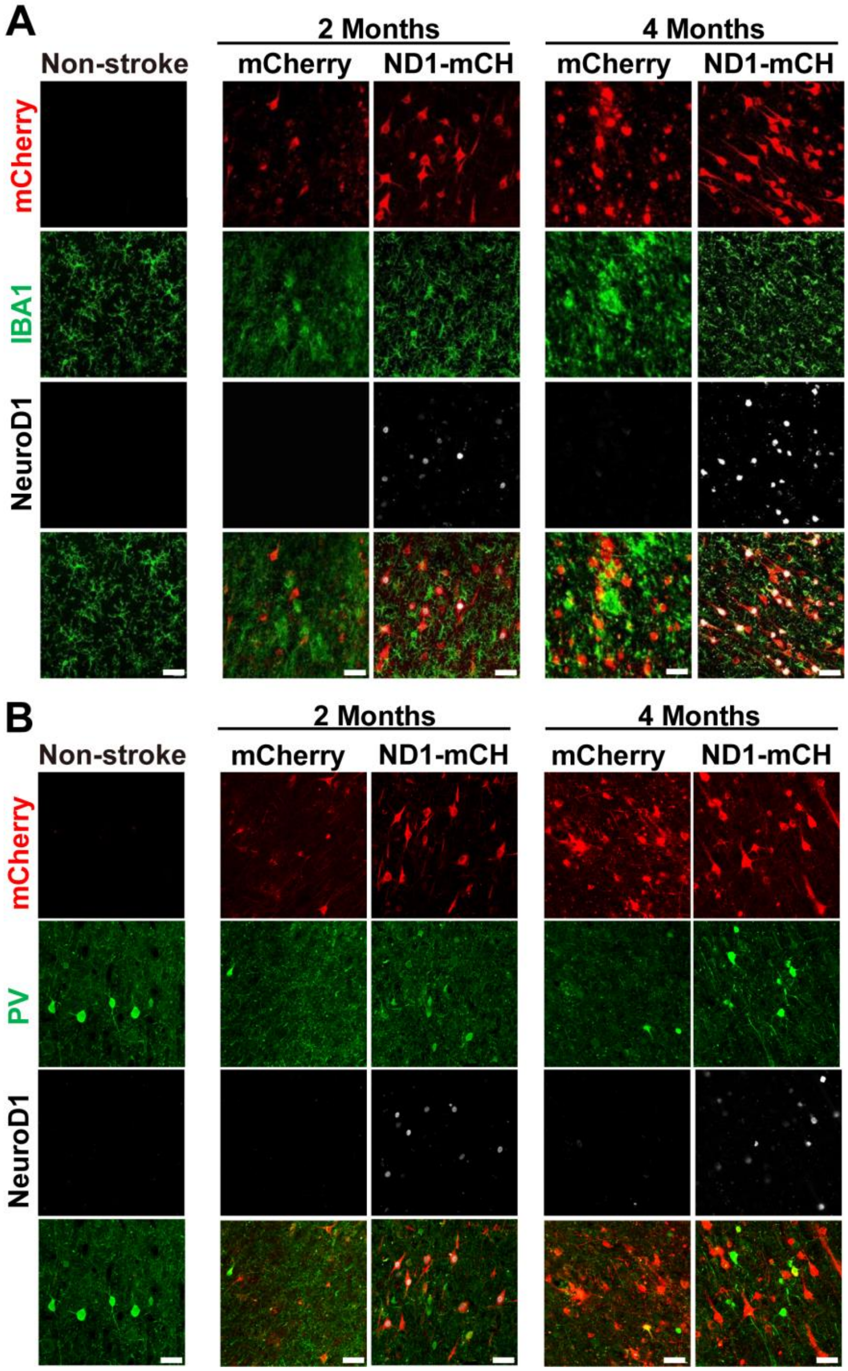


**Figure S5. NeuroD1-treatment reduces reactive microglia and promotes PV+ interneuron survival.**

**(A)** Representative images showing mCherry (red), Iba1 (green) and NeuroD1 (white) expression pattern in non-stroke cortex (left column) or ischemic cortex (right 4 columns) with virus injection at 10 days post stroke. Note that microglia (Iba1) showed less reactive morphology in NeuroD1-infected areas. Scar bars, 50 µm.

**(B)** Representative images showing mCherry (red), PV (green) and NeuroD1 (white) expression pattern in non-stroke cortex (left column) or ischemic cortex (right 4 columns). AAV infection at 10 days post stroke. Note a significant rescue of PV+ interneurons in NeuroD1-infected areas. Scar bars, 50 μm.

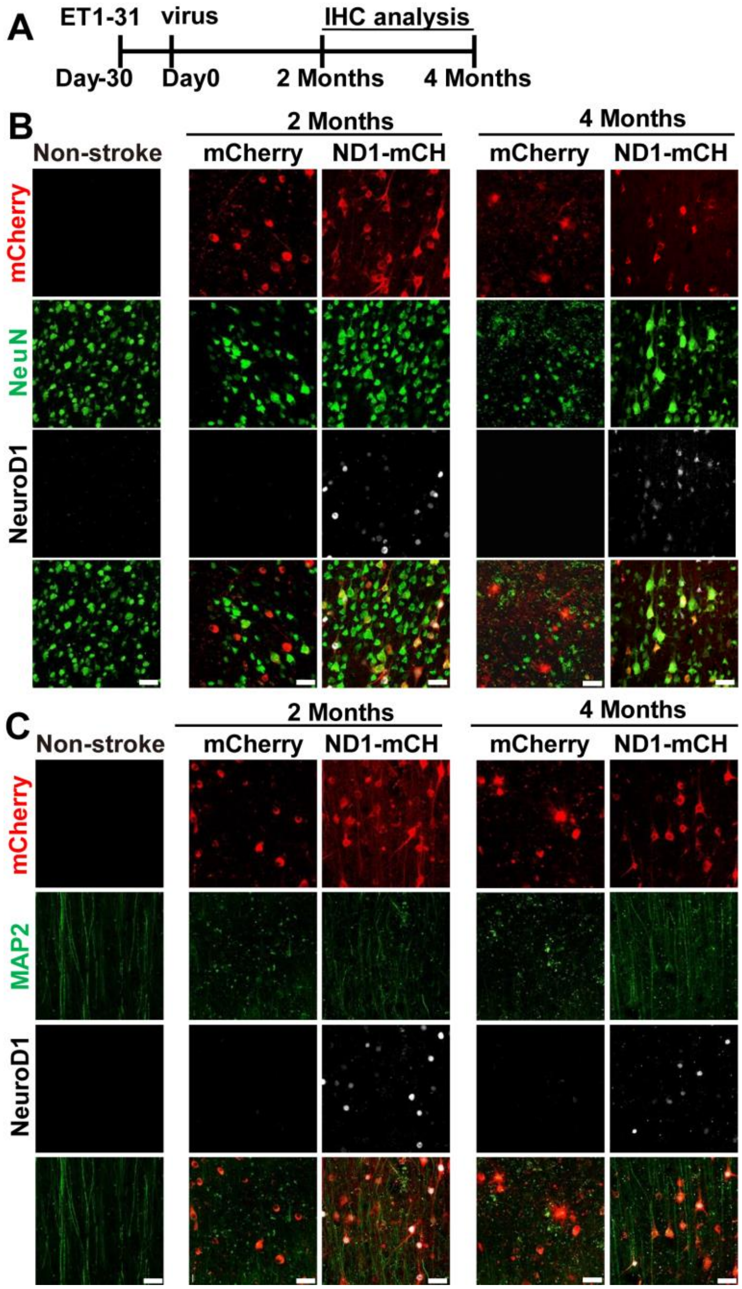


**Figure S6. NeuroD1-treatment at 30 days after ischemic stroke still rescued neuronal density in monkey cortex.**

**(A)** Timeline showing experimental design.

**(B)** Representative images showing mCherry (red), NeuN (green) and NeuroD1 (white) expression pattern in non-stroke cortex (left column) and ischemic cortex (right 4 columns) with virus infection at 30 days post stroke. Scar bars, 20 μm.

**(C)** Representative images showing mCherry (red), MAP2 (green) and NeuroD1 (white) expression pattern without (left column) or with stroke (right 4 columns). Scar bars, 20 μm.

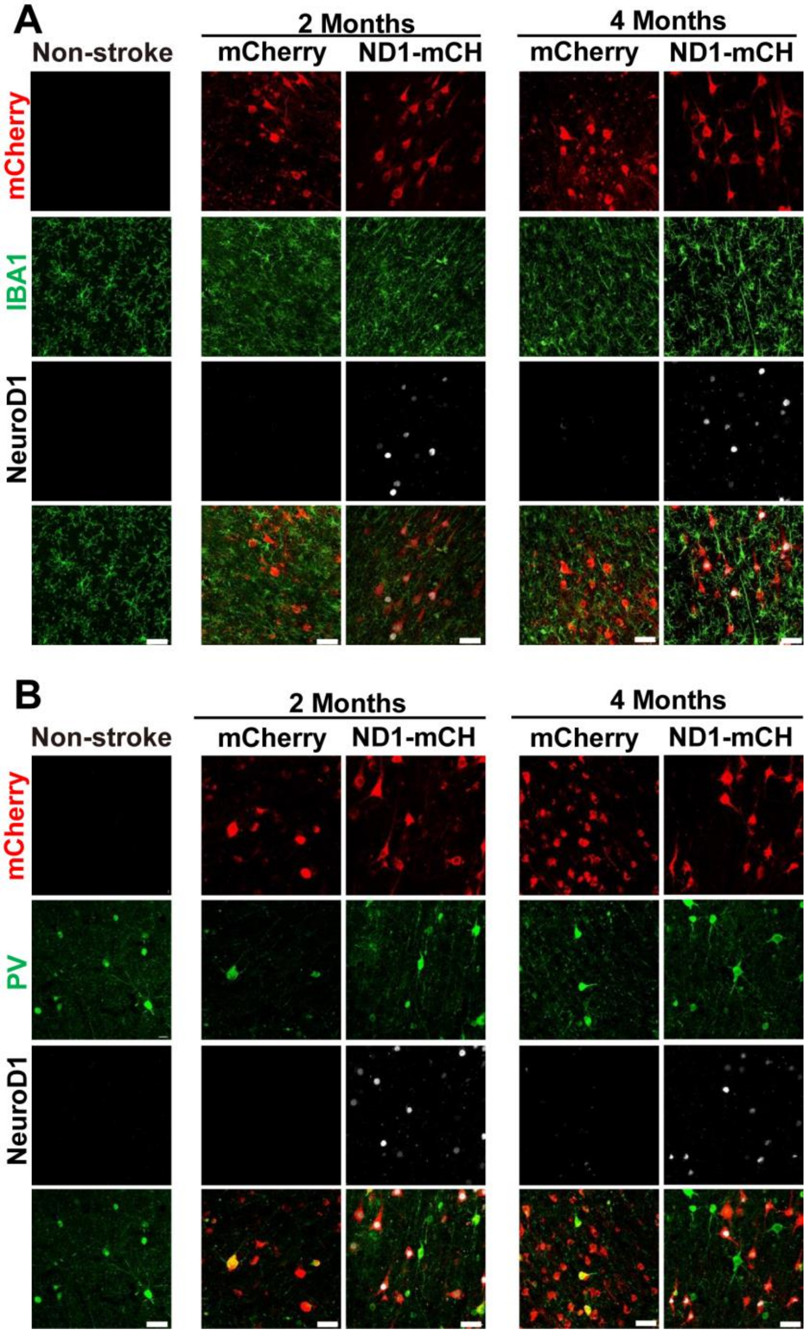


**Figure S7. NeuroD1-treatment at 30 days after ischemic stroke reduces reactive microglia and promotes PV+ neuron survival.**

**(A)** Representative images showing mCherry (red), Iba1 (green) and NeuroD1 (white) expression pattern in non-stroke cortex (left column) and ischemic cortex (right 4 columns). Virus infection at 30 days post stroke. Scar bars, 50 μm.

**(B)** Representative images showing mCherry (red), PV (green) and NeuroD1 (white) expression pattern in non-stroke cortex (left column) and ischemic cortex (right 4 columns) with virus infection at 30 days post stroke. Note that PV neurons were rescued in NeuroD1-infected areas. Scar bars, 50 μm.
